# A conserved ESCRT-II-like protein participates in the biogenesis and maintenance of thylakoid membranes

**DOI:** 10.1101/2023.10.10.561251

**Authors:** Irem Yilmazer, Pamela Vetrano, Simona Eicke, Melanie R. Abt, Eleonora Traverso, Tomas Morosinotto, Samuel C. Zeeman, Silvia Ramundo, Mayank Sharma

**Affiliations:** Institute of Molecular Plant Biology, Department of Biology, ETH Zürich, Zürich, Switzerland; Gregor Mendel Institute of Molecular Plant Biology, Austrian Academy of Sciences, Vienna BioCenter (VBC), Vienna, Austria; Department of Biology, University of Padova, Padova, Italy

**Keywords:** Photosynthesis, Thylakoid Biogenesis, ESCRT, Chloroplast, Protein-Protein Interaction

## Abstract

Thylakoids are membrane-bound compartments located in cyanobacteria and chloroplasts of plants and algae. They play an indispensable role in the light-driven reactions that enable photosynthetic organisms to convert water and carbon dioxide into oxygen and sugars. The biogenesis and maintenance of thylakoid membranes is a critical yet underappreciated area of research. One of the few known critical regulators of this process, VIPP1 (Vesicle-Inducing Protein in Plastids 1), was recently shown to be structurally similar to ESCRT-III proteins — the first evidence for ESCRT-like (Endosomal Sorting Complex Required for Transport) machinery in chloroplasts. Here, we used an affinity purification approach in two distantly related photosynthetic eukaryotes, the green alga *Chlamydomonas reinhardtii* and the plant *Arabidopsis thaliana*, to discover proteins that interact with VIPP1. Among several newly identified proteins, we focused on a highly conserved but uncharacterized protein (VIPP1-Associated protein 1, VIA1) that robustly interacts with VIPP1 in both systems. VIA1 is predicted to contain a winged-helix domain, a characteristic feature of ESCRT-II proteins that mediates the interaction with ESCRT-III proteins. The absence of VIA1 causes thylakoid swelling upon exposure to high light in Chlamydomonas and defective thylakoid biogenesis in the newly emerging leaf tissue in Arabidopsis, thereby delaying chloroplast development in this tissue. We propose that VIA1 is part of a previously unrecognized chloroplast ESCRT-like system that plays a critical role in forming, remodeling, and repairing photosynthetic membranes.

**Significance Statement:** Thylakoid membranes are essential for photosynthesis, yet their biogenesis and maintenance are poorly understood. Of the few known proteins involved in these processes, VIPP1 stands out due to its similarity to ESCRT-III, an integral component of the ESCRT machinery that is responsible for membrane remodeling and trafficking in the cytoplasm of eukaryotes. Here we report the discovery of VIA1, a conserved protein that interacts with VIPP1 and participates in thylakoid biogenesis and remodeling in two distantly related photosynthetic organisms. Because VIA1 contains a predicted winged-helix domain, a hallmark feature of ESCRT-II proteins that mediates the interaction with ESCRT-III proteins, our data support the hypothesis that universal, mechanistic principles govern membrane remodeling across all living organisms.

## Introduction

Photosynthesis, which is essential for much of life on Earth, is dependent on the extensive network of highly dynamic and specialized membranes known as thylakoids. The thylakoid membrane system presumably predates the origin of chloroplasts, as it is also present in cyanobacteria. While thylakoid lipid composition has remained highly conserved through evolution (1), the spatial organization of the membranes has undergone significant change over time (2, 3) suggesting that the biogenic process has also evolved. The complexity of the thylakoid membrane biogenesis —which involves the coordinated synthesis, assembly, and integration of lipids, proteins, and pigments— and its susceptibility to external stress stimuli have impeded efforts to elucidate this vital process. In fact, over the past twenty years, only a few of the molecular components directly involved in the biogenesis and maintenance of thylakoid membranes have been discovered (4–10).

One of the most extensively investigated of these factors is <u>V</u>esicle Inducing <u>P</u>rotein in <u>P</u>lastid 1 (VIPP1), a vital, conserved protein found in nearly all oxygenic photosynthetic organisms. Several studies have pointed to VIPP1’s involvement in thylakoid biogenesis (4, 11–13) as well as in chloroplast envelope and thylakoid membrane maintenance (14–17). Most notably, deletion of the VIPP1 gene in the land plant *Arabidopsis thaliana* and *Synechocystis* sp. PCC6803, was shown to induce severe abnormalities in their thylakoid membrane architecture (4, 11, 15). Another study in the unicellular green alga *Chlamydomonas reinhardtii* demonstrated that downregulation of VIPP1 via RNA interference causes light-induced thylakoid swelling (14). A similar phenotype was also observed in Arabidopsis chloroplasts upon knockdown of VIPP1 (15) and in Synechocystis cells expressing a mutant variant of VIPP1 (18).

Recent groundbreaking studies of the cyanobacterial VIPP1 revealed that this protein has a common evolutionary origin and shares functional and structural similarities with the <u>E</u>ndosomal <u>S</u>orting <u>C</u>omplex <u>R</u>equired for <u>T</u>ransport (ESCRT)-III components of eukaryotic cells (18, 19). The ESCRT system plays crucial roles in endosomal and non-endosomal processes related to protein sorting and membrane remodeling and is composed of ESCRT-0, -I, -II, and -III protein complexes, as well as accessory proteins (20). The proteins of this highly conserved pathway were considered a hallmark of eukaryotic cells for many years. However, the discovery of ESCRT-III homologs in archaea, cyanobacteria, and chloroplasts suggests the existence of an ancient ESCRT-like pathway (18, 19, 21–23). It therefore stands to reason that this pathway likely encompasses additional conserved components beside VIPP1, although their functional impact may differ, much like VIPP1 itself.

To discover some of these additional components, we conducted co-immunoprecipitation studies in *A. thaliana* and *C. reinhardtii* to find proteins that interact with VIPP1. We identified a few known and several uncharacterized VIPP1-interacting proteins. Among the uncharacterized proteins was a protein that shares structural similarities with components of the eukaryotic ESCRT-II complex. This protein, which we named <u>VI</u>PP1-<u>A</u>ssociated protein 1 (VIA1), is conserved across cyanobacteria, green algae, and land plants. Our analysis of VIA1, presented herein, points to it being a member of a conserved molecular machine involved in chloroplast membrane biogenesis and maintenance.

## Results

### 1. Identifying the VIPP1-interactome in Arabidopsis

To determine the protein-interaction network of VIPP1, leaf tissues from three independent transgenic Arabidopsis lines expressing VIPP1-YFP were used as the starting material to co-immunoprecipitate VIPP1-YFP and associated proteins with the help of GFP-trap magnetic beads. This was followed by on-bead protein digestion and subsequent mass spectrometry analysis of the released peptides. Wild type plants (WT Col-0) and plants expressing plastid-stroma targeted GFP (24) were used as controls to identify potentially false-positive interactions. In total, this affinity purification coupled to mass spectrometry (AP-MS) approach led to the detection of peptides corresponding to 369 different proteins, which were quantified according to peptide intensities across the samples. The bait protein, VIPP1-YFP, ranked first, with the highest count of identified peptides. Eight additional proteins emerged as potential VIPP1 interacting partners (Figure 1). These proteins consistently appeared across all biological replicates but were absent or substantially reduced in all control samples. Moreover, all eight selected proteins were previously detected in chloroplast proteomic studies and/or carried a predicted chloroplast transit peptide (25). Two of these eight proteins, <u>H</u>eat <u>S</u>hock 70 kDa <u>P</u>rotein-7 (HSP70-7) and chaperone Dna-J domain-containing protein C73 (DnaJ-C73), share homology with Chlamydomonas HSP70B and CDJ2, respectively. This chaperone pair was previously reported to interact with VIPP1 and regulate its oligomerization state (26, 27). Their presence provides strong validation of our AP-MS method. Of the six remaining proteins, the second most abundant after VIPP1 was At5g03900, the homolog of CPLD50/VPL3, an uncharacterized protein that was prominently featured in a recent VIPP1-proximity labelling study in Chlamydomonas, together with HSP70B and CDJ2 (28). Two of the other VIPP1-interacting partners identified in this analysis are uncharacterized proteins containing predicted transmembrane domains. The first belongs to the <u>P</u>entatricope<u>p</u>tide <u>R</u>epeat (PPR) protein family (29), a large and diverse family of proteins characterized by the presence of a repetitive 35 amino acid motif. The second is a small,10 kDa protein that was detected within the chloroplast inner envelope in a prior proteomics study (30). Of the three remaining VIPP1-interacting proteins, one is a verified chloroplast membrane protein, <u>F</u>luctuating <u>L</u>ight <u>A</u>cclimating <u>P</u>rotein (FLAP1), involved in acclimation response to variable light conditions (31). The other two are predicted components of chloroplast enzyme complexes: a small subunit of the <u>A</u>cetolactate <u>S</u>ynthase protein complex (ALS-2) and a plastid-encoded NAD(P)H-quinone oxidoreductase subunit I (NADH-I). While some of these proteins have been previously studied, their roles in the context of VIPP1 function are yet to be determined.

**Figure 1.**
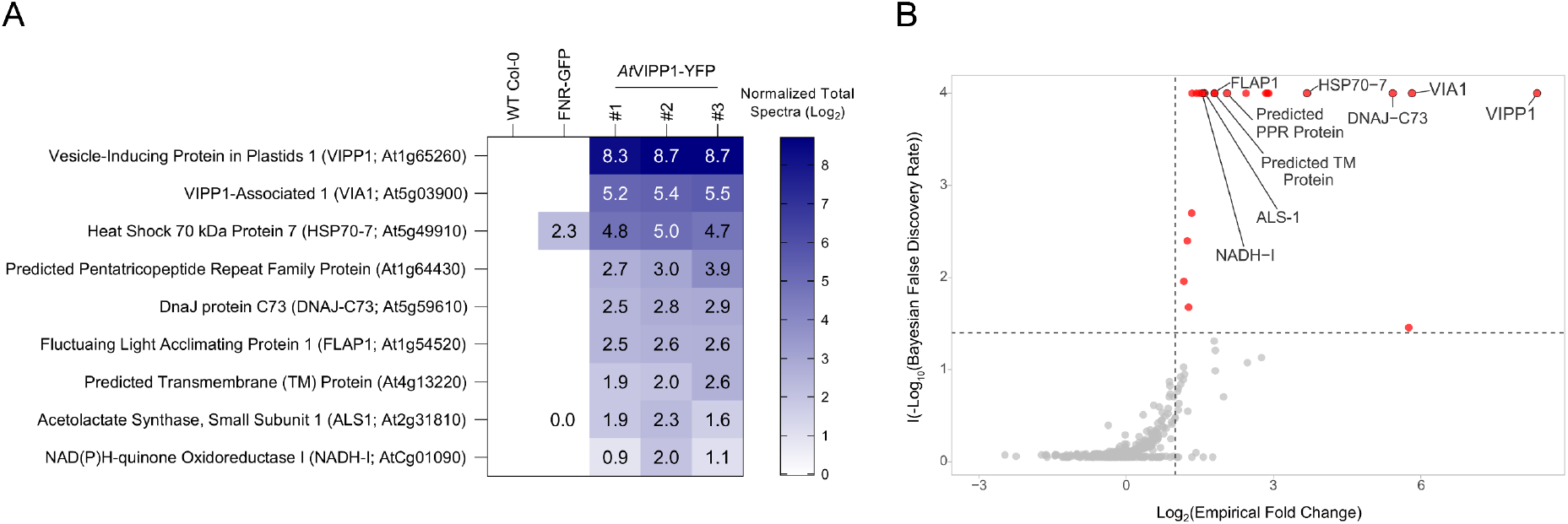
Identification of VIPP1 Interacting Partners in *Arabidopsis thaliana*. **(A)** Heatmap representing a group of proteins identified in an AP-MS experiment conducted with Arabidopsis leaf total protein lysates expressing VIPP1-YFP. These proteins showed negligible or undetectable levels in control samples (WT Col-0 and FNR-GFP) (Suppl. File 2). Scaffold proteomics software was employed to analyze the identified peptides, and a quantitative value representing the identified peptide (normalized total spectra) is displayed within each cell of the heatmap. The values are Log_2_-transformed. The heatmap was generated using GraphPad Prism software. **(B)** Volcano plot showing −log_10_ transformed Bayesian False Discovery Rate (BFDR) as a function of log_2_ transformed effect size. The dotted lines show −log_10_ transformed BFDR threshold of 0.035, and the log_2_ transformed empirical fold change of 2. Proteins that lie above this threshold are highlighted in red. SAINTexpress software was utilized to score the interaction between bait and prey proteins.

### 2. VIA1 interacts with VIPP1 and co-localizes in punctate structures

The uncharacterized protein encoded by At5g03900 caught our attention as it was the second most enriched protein in our AP-MS experiment. Moreover, this protein, which we named VIPP1-Associated protein 1 (VIA1), contains three predicted transmembrane domains and has previously been detected in the inner membrane fractions of chloroplast envelope in several proteomic analyses (30, 32, 33). To confirm the interaction between VIA1 and VIPP1, we employed a reverse IP approach. Yellow fluorescent protein (YFP)-tagged VIA1 was used as a bait and transiently expressed in leaf epidermis cells of *Nicotiana benthamiana* together with FLAG-tagged VIPP1. In this experiment, another YFP-tagged chloroplast protein (cpFeS, involved in iron-sulphur cluster biogenesis) served as a negative control bait. In all replicates, VIPP1-FLAG co-precipitated with VIA1-YFP as shown by an immunoblot analysis (Figure 2A). In contrast, no interaction between VIPP1-FLAG and the negative control bait could be detected. Moreover, the binding of VIPP1-FLAG to the empty GFP-trap magnetic beads was negligible. Collectively, these findings strongly suggest that VIA1 interacts with VIPP1 in a selective manner.

**Figure 2.**
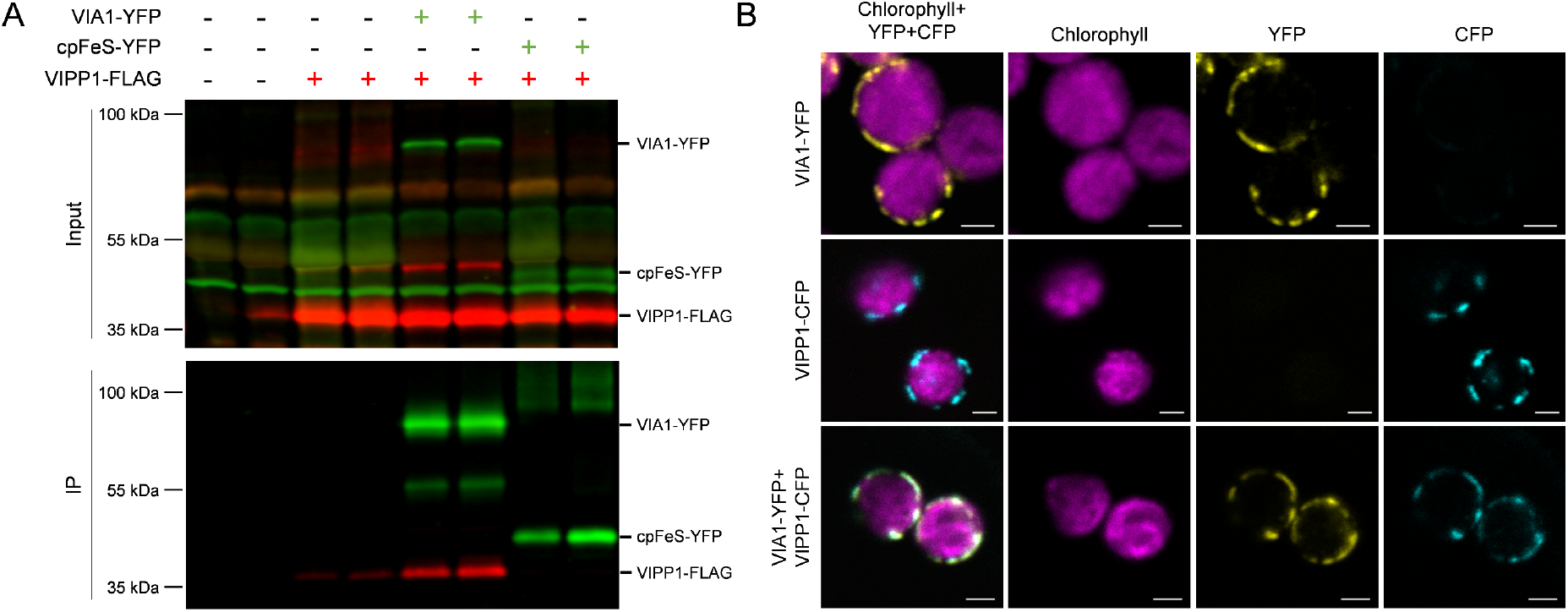
VIA1 Interacts with VIPP1 and Colocalizes in Distinct Puncta in Chloroplasts. **(A)** Co-immunoprecipitation experiment involving transiently overexpressed VIA1-YFP and VIPP1-FLAG in *N. benthamiana* leaf tissue. Proteins samples were separated via SDS-PAGE, transferred onto a PVDF membrane, and detected with rabbit anti-GFP and mouse anti-FLAG primary antibodies. Visualization was achieved using secondary antibodies labeled with distinct infrared (IR) fluorescent dyes specific for rabbit IgG (green) and mouse IgG (red). **(B)** Confocal laser scanning microscopy analysis of chloroplasts in lower epidermis cells of *N. benthamiana* leaf tissue expressing VIA1-YFP, VIPP1-CFP, or both. Chlorophyll signals are false-colored magenta, YFP signals yellow, and CFP cyan. Scale bar represents 2 µm.

Next, we reasoned that if VIA1 interacts with VIPP1, both proteins should co-localize in plant cell chloroplasts. To test this, we transiently co-expressed VIPP1 fused with a cyan fluorescent protein (CFP) together with VIA1-GFP, and determined the subcellular localization of the two proteins using confocal laser scanning microscopy (CLSM). We found that VIPP1 and VIA1 co-localized to punctate structures primarily located at the periphery of the chloroplast (Figure 2B, Figure S1). These results align with the previously reported chloroplast envelope localization of VIA1 and provide further evidence for the interaction between VIA1 and VIPP1 *in vivo*.

### 3. VIA1 is evolutionarily conserved

It has been hypothesized that VIPP1 evolved via duplication of the bacterial <u>P</u>hage <u>S</u>hock <u>P</u>rotein <u>A</u> (PspA) gene and is associated with the origin of the thylakoid membrane system (11). VIPP1 is conserved in all photosynthetic organisms harboring a thylakoid membrane system studied to date (13). To determine whether VIA1 is likewise evolutionarily conserved, we identified its potential homologs using NCBI-BLAST against the refseq_proteins dataset and conducted phylogenetic analyses using the maximum likelihood method. This revealed the presence of VIA1 homology across land plants, algae, and cyanobacteria, including members of the *Gloeobacter* lineage, which lack thylakoids (Figure 3, Suppl. File 3). While most of the thylakoid-lacking cyanobacteria do not contain a *bona fide* VIPP1 protein, the presence of VIA1 in these organisms suggests an alternate function for this protein.

**Figure 3.**
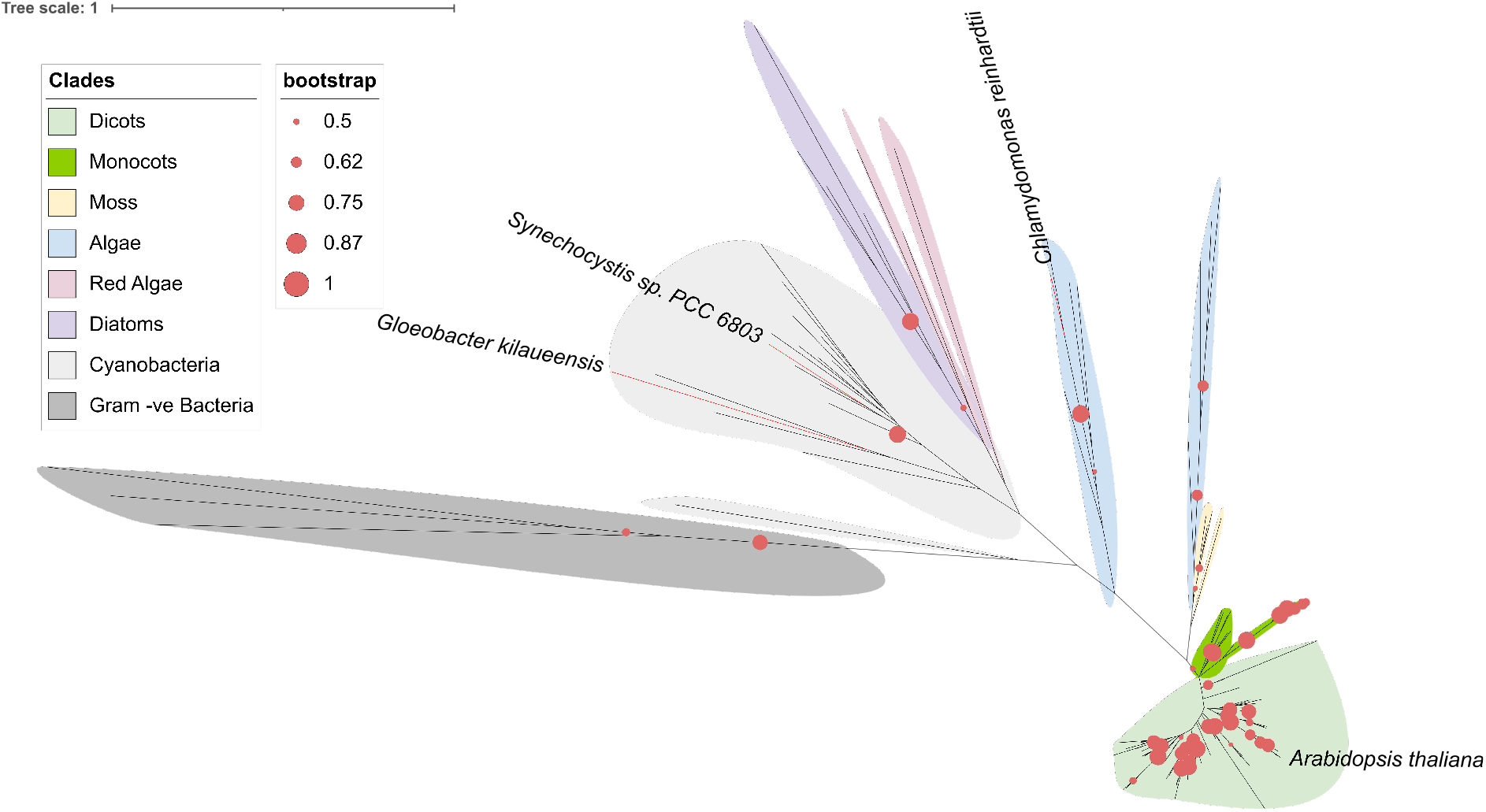
Evolutionary Conservation of VIA1. A maximum likelihood phylogeny of VIA1 sequences from 140 different species was computed using the LG+G+F model for protein evolution with MEGA7 software. One thousand bootstrap replications were performed and an unrooted tree with scaled branches is presented. Each branch corresponds to a specific species, and branches are organized into clades, color-coded as indicated in the legend at the top-left corner. Bootstrap values of more than 50% are depicted as small pink dots at the base of each branch. The bootstrap range is indicated in the legend. The phylogenetic tree was annotated using iTOL (Letunic and Bork, 2021) web-based software. Branches corresponding to the species *Gloeobacter kilaueensis*, *Synechocystis* sp. PCC6803, *Chlamydomonas reinhardtii*, and *Arabidopsis thaliana* are colored red. See Suppl. File 3 for a fully annotated tree in rectangular format.

### 4. VIA1 (CPLD50) interacts with VIPP1 in *C. reinhardtii*

To examine if VIA1’s interaction with VIPP1 is conserved in other photosynthetic organisms, we took an orthogonal AP-MS approach using *C. reinhardtii*, where VIPP1 has been extensively characterized. Cell lysates from a wild-type (WT) Chlamydomonas strain were incubated either with Dynabeads protein-A coupled with a polyclonal anti-VIPP1 antibody or with non-coupled Dynabeads (negative control). A total of 491 proteins were identified across all three technical replicates, with VIPP1 being the most abundant, confirming the specificity of the anti-VIPP1-antibody conjugated beads (Figure 4). Heat Shock Protein 70B (HSP70B) and chloroplast DNAJ-like chaperone 2 (CDJ2) were abundantly present, as in Arabidopsis. We also detected VIPP2, a VIPP1 paralog known to interact with VIPP1 (34), alongside a series of other uncharacterized proteins, some of which (namely, VPL1-6 and VPL10) were recently identified in a VIPP1 proximity-labeling experiment (28). Importantly, peptides corresponding to CPLD50 (35), the ortholog of Arabidopsis VIA1 (hereafter “*Cr*VIA1”), were significantly enriched in all samples except the control, confirming that the interaction between VIA1 and VIPP1 is conserved in *C. reinhardtii*. This list also included other proteins akin to those identified in the *A. thaliana* VIPP1 AP-MS analysis; for example, CPLD42 (an ortholog of *At*FLAP1), Cre14.g611950, a putative ortholog of the PPR protein (At1g64430), and Cre16.g687200, an ortholog of the transmembrane protein (At4g13220). The identification of this comparable set of VIPP1 interacting proteins in both Chlamydomonas and Arabidopsis strengthens our conclusion that these proteins are *bona-fide* VIPP1 interacting partners.

**Figure 4.**
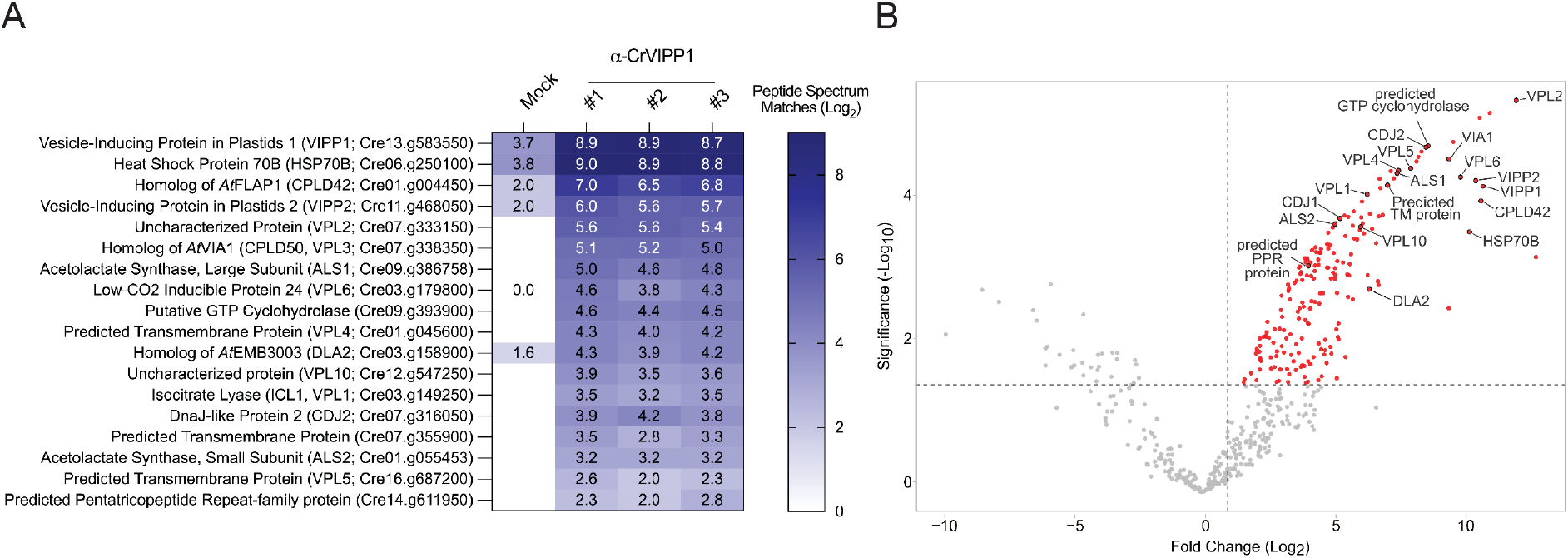
Identification of VIPP1 Interacting Partners in *C. reinhardtii*. **(A)** A heatmap showcasing proteins consistently identified across VIPP1 co-immunoprecipitation experiments in three biological replicates. These proteins showed negligible or undetectable levels in control samples (i.e., wild-type lysates incubated with uncoupled Protein A Dynabeads) (Suppl. File 4). The values shown within each cell of the heatmap represent the Peptide Spectrum Matches (PSMs) for each protein, serving as an indicator of their relative abundance. The heatmap was generated using GraphPad Prism software. **(B)** Volcano plot displaying log_2_ transformed fold change (FC) and -log_10_ transformed p-values for each confidently quantified protein. Proteins significantly enriched in the VIPP1 co-immunoprecipitation are highlighted in red. The labels within the plot correspond to the proteins of interest in the accompanying heatmap.

### 5. VIA1 is structurally similar to ESCRT-II proteins and is predicted to interact with VIPP1 via its winged-helix domain

To understand the molecular basis of the interaction between VIPP1 and VIA1, we carried out structural predictions using Alphafold2. Despite the relatively low sequence identity (38%) between CrVIA1 and AtVIA1, the two proteins appear to exhibit a notable structural similarity (Figure 5). This is supported by a root mean square deviation (RMSD) of 3.07 and a template modeling (TM) score of 0.72, as determined through Pairwise Structure Alignment, an online tool from RCSB-PDB (36) (Figure S2). Since no conserved domain could be identified through primary amino acid sequence analysis, we used the BackPhyre tool to search for proteins with structural similarities (37). Among the top hits was an ESCRT-II protein, Snf8, from *Saccharomyces cerevisiae*. Our subsequent analysis using Swiss Model and Phyre2 revealed that VIA1 carries a winged-helix (WH) domain at its N-terminus similar to the WH domains of ESCRT-II proteins (Figure 5). Additionally, opposite of this WH domain is a helix-turn-helix motif that folds in a WH domain-like topology.

**Figure 5.**
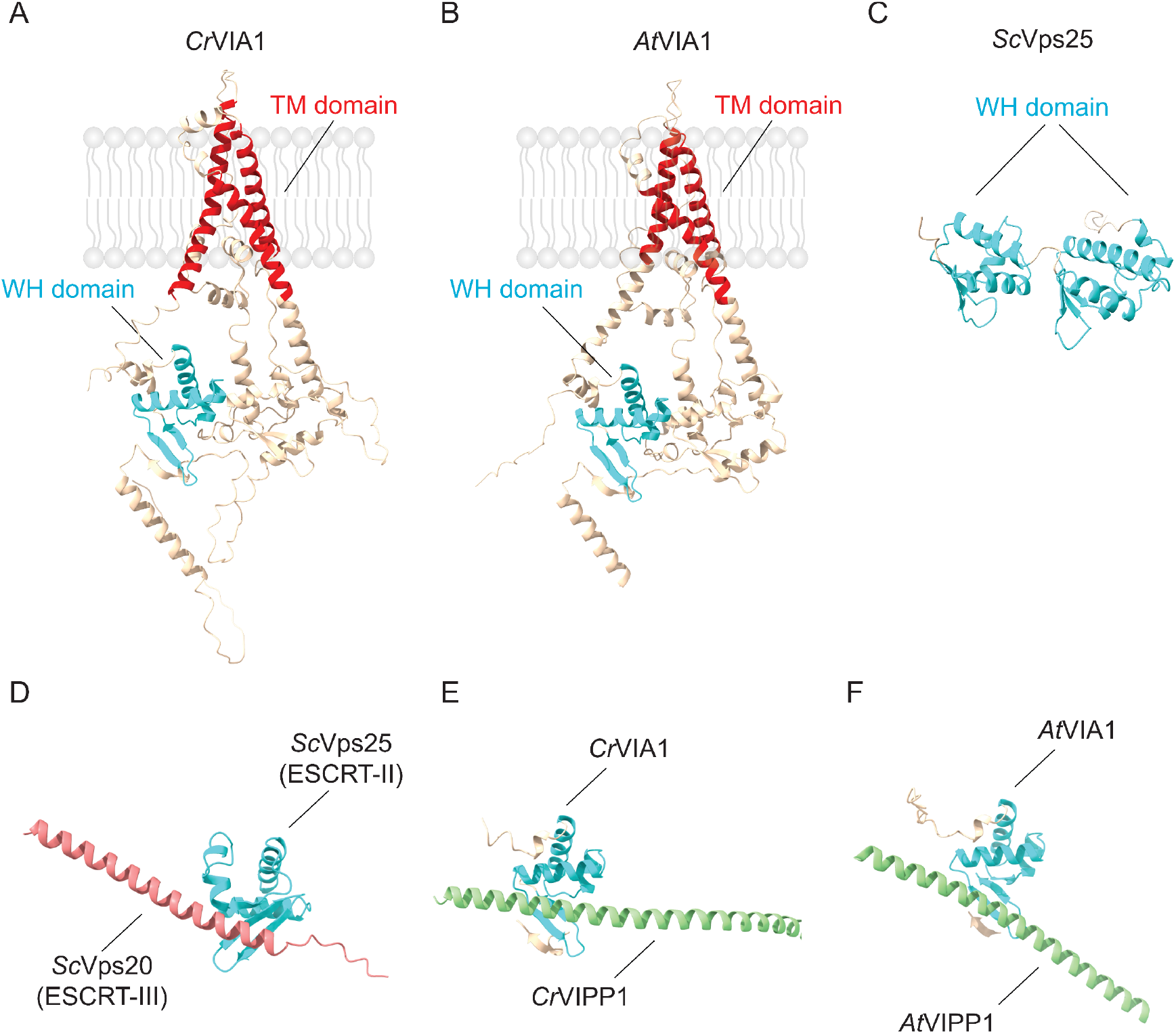
Structural Analysis of VIA1 and its Interaction with VIPP1. Alphafold2 structural predictions of VIA1 from *C. reinhardtii* (panel **A**) and *A. thaliana* (panel **B**). The winged helix (WH) domains are colored in cyan, and transmembrane (TM) domains in red. (**C**) Experimentally determined structure of Vps25 from *S. cerevisiae*. (**D**) The interaction between *Sc*Vps25 (ESCRT-II subunit) and *Sc*Vps20 (ESCRT-III subunit) is illustrated. This interaction was experimentally determined (38) and takes place at the interface between the first α-helix of *Sc*Vps20 (in pink) and the hydrophobic pocket within the WH domain of *Sc*Vps25 (in cyan). Alphafold2-predicted interaction of VIA1 (in cyan) and VIPP1 (in green) in Chlamydomonas (panel **E**) and Arabidopsis (panel **F**).

In eukaryotes, the WH domain is present in ESCRT-II subunits, such as Vps25, Vps36, and Vps22 (also called Snf8) and is critical for interaction with components of the ESCRT-III complex (39). This ESCRT-II-ESCRT-III interaction involves the formation of a hydrophobic surface patch by the ESCRT-II-WH domain upon interacting with the first α-1 helix of the ESCRT-III protein (38). To determine if this is also true for the interaction between VIPP1 and VIA1, we modeled the VIPP1-VIA1 interaction using Alphafold multimer (40) and compared it to that of two well-characterized ESCRT-III and ESCRT-II subunits, Vps20 and Vps25, respectively. We observed a striking similarity between the experimentally determined ESCRT-II and ESCRTIII subunits’ interface and the predicted VIPP1-VIA1 interface (Figure 5E, F). This predicted similarity lends support to the hypothesis that VIA1 and VIPP1 function in chloroplasts analogously to the components of the cytosolic ESCRT machinery.

### 6. Loss of CrVIA1 leads to *C. reinhardtii* hypersensitivity to photooxidative stress

VIPP1 is essential for the formation of thylakoids and their maintenance in response to high light in Chlamydomonas. The ultrastructure of thylakoid membranes in *VIPP1* knockdown strains is significantly different from WT and the mutants are extremely sensitive to photooxidative stress (14). To determine if VIA1 is functionally related to VIPP1, we obtained two different Chlamydomonas *VIA1* insertional mutant lines, hereafter referred as *via1-1* and *via1-2*, generated during the Chlamydomonas Library Project (CLiP) (41). The mutagenic insertion in the *Cr*VIA1 locus in both strains was validated using PCR (Figure S3) and backcross analyses (Figure S4). Moreover, qRT-PCR analysis suggested that VIA1 expression is strongly reduced in both *via1-1* and *via1-2* (Figure 6A, B).

**Figure 6.**
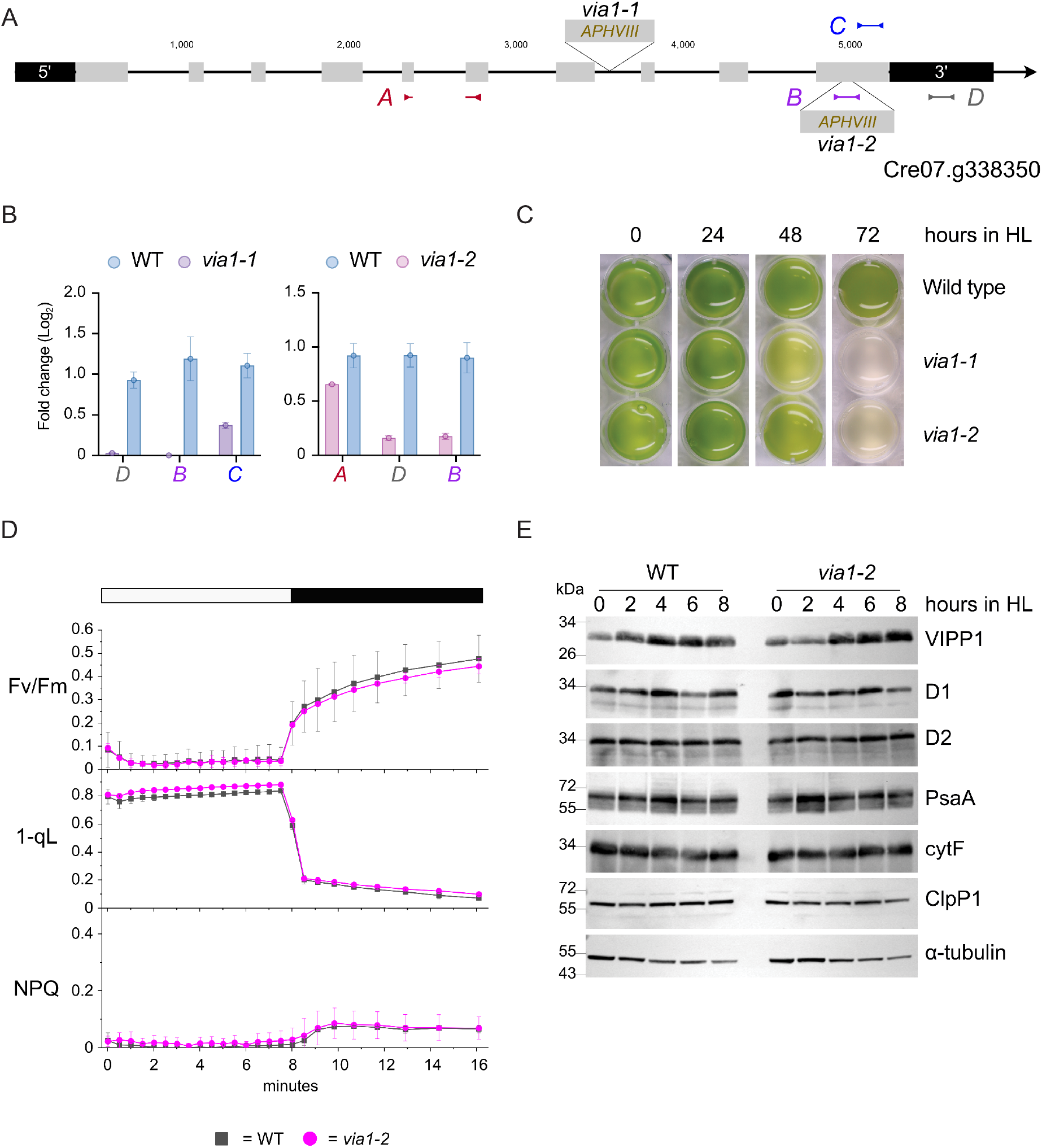
Validation and Phenotypic Characterization of Chlamydomonas Mutants Lacking VIA1. (**A**) Schematic representation of *CrVIA1*, with exons depicted in gray and 5’ and 3’ untranslated regions in black. The position of the mutagenic cassette carrying the paromomycin resistance gene (*APHVIII*) in *via1-1* and *via1-2* is indicated. The colored lines represent three *VIA1* regions selected for the quantitative RT-PCR analyses presented in panel B. **(B)** *VIA1* expression levels in *via1-1* and *via1-2* strains. The x-axis labels are color-coded according to the scheme depicted in panel A. *GBLP* was used as a reference gene for normalization **(C)** Images of liquid TAP cultures of WT, *via1-1*, and *via1-2* strains at various time points during a 3-day high light (HL) treatment. One of three biological replicates is displayed. (D) Photosynthetic measurements in WT and *via1-2* cells using PAM fluorometry. Photosystem II quantum yield (Fv/Fm) was monitored during an 8-minute exposure to actinic light at an intensity of 1000 µmol photons m^-2^ s^-1^ (white bar), followed by an 8-minute dark period (dark bar). The Plastoquinone redox state was assessed by analyzing the fluorescence parameters 1-qL and NPQ. Data are presented as means ± standard deviations from four independent measurements. (E) Immunoblot analysis of cell extracts following HL treatment for 0, 2, 4, 6, and 8 h. The blots were probed with antibodies against VIPP1, D1, D2, PsaA, cytF, ClpP1, and α-tubulin.

When grown under standard light conditions (100 µmol photons m^-2^ s^-1^), *via1-1* and *via1-2* were indistinguishable from WT, as illustrated by measurements of various photosynthetic parameters such as photosystem II efficiency, non-photochemical quenching, and plastoquinone (PQ) redox state (Figure 6D, Figure S5). However, both *VIA1* mutants were more sensitive to photooxidative stress than WT when exposed to high light (HL, ∼800 µmol photons m^-2^ s^-1^) (Figure 6C). In particular, chlorophyll content was significantly more reduced in mutants than in WT after 2 days (Figure S6) and photobleaching occurred earlier in the mutants than in WT.

Next, we utilized transmission electron microscopy (TEM) to monitor chloroplast membrane ultrastructure in both WT and *via1* cell lines at 2-h intervals following HL exposure. Thylakoids were extremely swollen in *via1-2* cells as early as 4 h after the start of the HL treatment (Figure 7B), whereas thylakoids in WT cells remained unaffected until 6 h in HL (Figure 7A). During the same time-course, we also measured expression of the photosynthetic machinery components PsaA (a core subunit of photosystem I), D1 and D2 (core proteins of photosystem II), and cytochrome F (the major subunit of the cytochrome b_6_f complex) (Figure 6E). We did not detect any significant differences in the steady-state levels of these proteins. We conclude that the increased photooxidative sensitivity observed in *via1-*2 under HL exposure does not stem from impaired assembly or function of the photosynthetic protein complexes. Rather, our data indicate that VIA1 is required for preserving thylakoid membrane integrity in response to chloroplast membrane damaging factors such as photooxidative stress.

**Figure 7.**
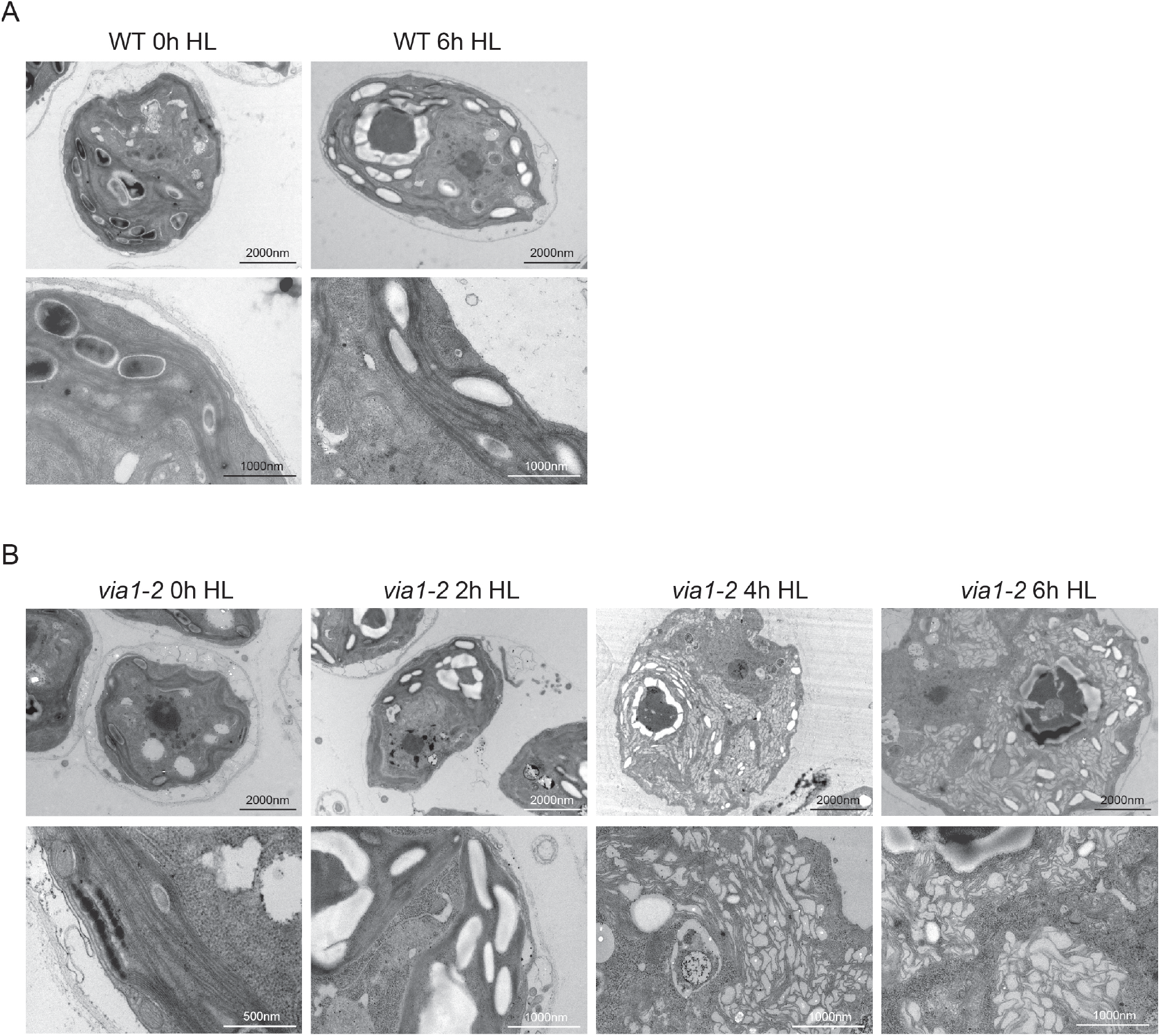
Enhanced Thylakoid Swelling in Chlamydomonas Chloroplasts Lacking VIA1 upon High Light (HL) Exposure. Transmission electron microscopy (TEM) images of cells exposed to HL (∼800 µmol photons m^-2^ s^-1^) for varying durations. **(A)** Representative TEM images of WT cells captured at 0- and 6-hours following HL exposure. **(B)** Representative TEM images of *via1*-2 cells captured at 0-, 2-, 4-, and 6-hours following HL exposure.

### 7. Loss of VIA1 causes defects in chloroplasts of emerging *A. thaliana* leaf tissue

To investigate if lack of VIA1 causes a similar alteration in thylakoid membrane structure in Arabidopsis, we obtained a *VIA1* insertion mutant line, SALK_057879C, and isolated plants homozygous for the T-DNA insertion using PCR-based genotyping (Figure 8A, B). The absence of VIA1 transcripts in this line was confirmed using semi-quantitative RT-PCR (Figure 8C) and the absence of VIA1 protein was tested using a co-immunoprecipitation experiment after introduction of VIPP1-YFP in the *via1* background (Figure S7). The *via1* plants appeared similar to WT Col-0 plants except that the basal area of newly emerging leaf tissue, adjacent to the rosette axis, was pale green in color (Figure 8D). To verify that the observed phenotype is due to a VIA1 deficiency, we rescued *via1* plants by introducing a transgene coding for FLAG tagged VIA1 under the control of its native promoter (*via1/proVIA1:VIA1FLAG*). This transgene complemented the phenotypic defect of *via1* in independent lines, with newly emerging leaf tissue resembling WT Col-0.

**Figure 8.**
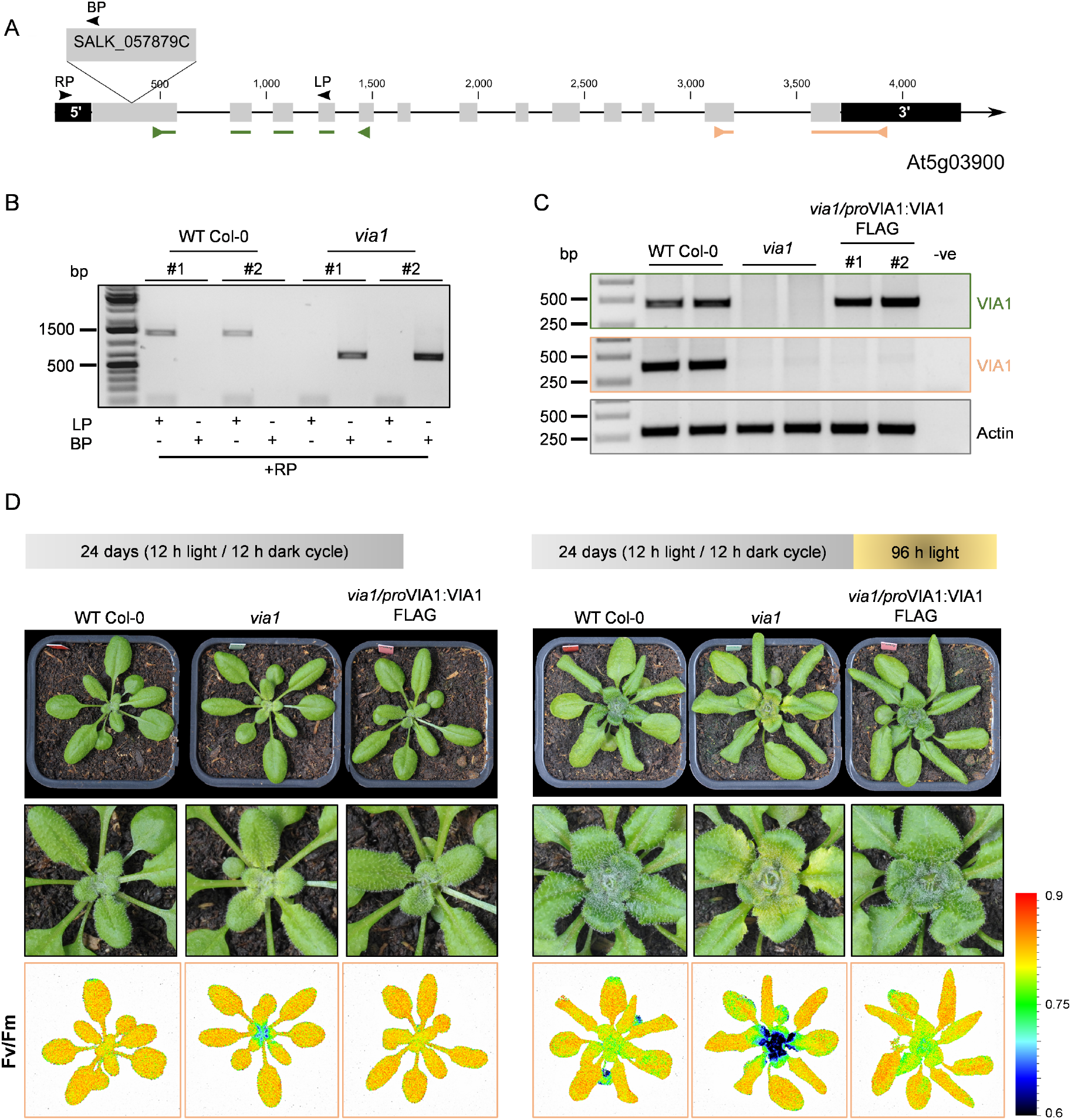
Functional Characterization of the *VIA1* Gene in Arabidopsis. **(A)** Schematic representation of the Arabidopsis *VIA1* gene indicating the position of the T-DNA insertion. Exons are depicted in gray, and 5’ and 3’ untranslated regions are shown in black. The colored lines represent *VIA1* regions selected for RT-PCR analysis, black arrows represent primer positions for genotyping **(B)** PCR-based genotyping of *via1* and wild type (WT Col-0) lines using T-DNA border (BP) and gene specific primers (LP and RP) as detailed in Table S1 **(C)** Semi-quantitative RT-PCR analysis of *VIA1* transcripts in WT Col-0, *via1,* and complementation lines. Amplification regions are color-coded according to panel A. **(D)** Photographs of 24-days-old plants (left) and with additional exposure to continuous light for 96 h (right). Depicted are one of three biological replications. Emerging leaf tissues *via1* plants display reduced photosynthetic efficiency. This reduction becomes more pronounced after an additional 96 h continuous light exposure. Images in the bottom panels represent the maximum quantum yield of PSII (Fv/Fm).

Using a closed FluorCam, we measured the maximum quantum yield of photosystem II (Fv/Fm) in all three lines (WT, *via1*, and *via1/proVIA1:VIA1FLAG*) after a short adaptation period in the dark. While the majority of the leaf tissue had a similar Fv/Fm ratio, the proximal part of newly emerging leaves (leaf base, i.e., the zone of extensive cellular proliferation) displayed a reduction in quantum yield of PSII solely in the *via1* line (Figure 8D). To investigate if this defect was more pronounced under conditions of constitutive photomorphogenesis, we transferred the plants to continuous light for 96 h and repeated the analysis. As anticipated, the region of pale green tissue at the base of newly emerging leaf tissue significantly expanded in *via1*, but not in WT Col-0, leading to a further reduction in the Fv/Fm ratio in that particular region. This phenotype was consistently observed throughout the initial developmental stages of plant growth, as evidenced by data collected from 14- and 28-day-old plants (Figure S8, S9). Since this phenotype was not observed in either the WT Col-0 or *via1/proVIA1:VIA1FLAG* lines, our findings indicate that the emergence of pale green tissue in *via1* is caused by the absence of VIA1 (Figure 8D, Figure S9).

Pale green leaf phenotypes often correlate with defective chloroplast development or homeostasis. To examine chloroplast architecture in *via1* leaf tissue, we carried out transmission electron micrography. In the green leaf tissue, the chloroplast ultrastructure was similar to that of WT Col-0 plants, with the exception of sporadic occurrences of *via1* chloroplasts with swollen thylakoid membranes (Figure S10, S11). In contrast, the thylakoid membrane network in chloroplasts from the pale green leaf tissue of *via1* was reduced (Figure 9A). Additionally, numerous vesicular structures accumulated within chloroplasts of *via1* (Figure S12), and the chloroplasts were smaller and distorted when compared to WT Col-0. To further characterize the morphology of chloroplasts in this region, we used serial block-face scanning electron microscopy (SBF-SEM). This method revealed that chloroplasts from WT Col-0 leaf tissue had a typical discoid shape, whereas those in leaf tissue of *via1* plants were distorted and irregular in shape (Figure 9B, Video S1, S2). The drastic decrease in thylakoid membrane network, along with the visible morphological abnormalities observed in plants lacking VIA1, highlight the importance of this newly discovered protein in thylakoid biogenesis and the maintenance of chloroplast structural integrity and functionality in newly formed leaf tissue.

**Figure 9.**
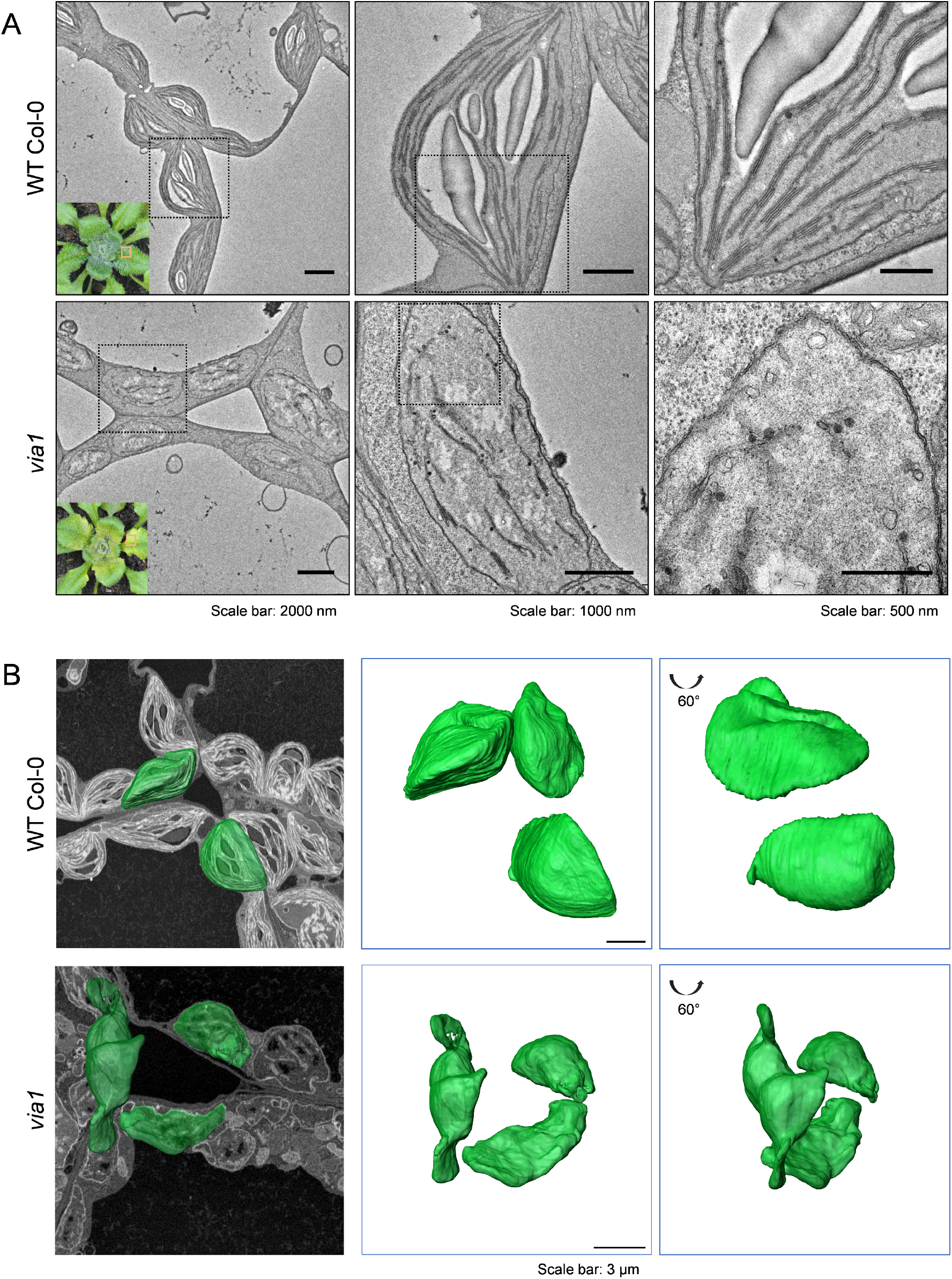
Abnormal Thylakoid Membranes Ultrastructure and Chloroplast Morphology in Arabidopsis *via1* mutant. **(A)** Transmission electron micrographs displaying chloroplasts from the highlighted region at the leaf base of 24-day-old wild-type and *via1* plants grown for an additional 96 h in continuous light. The micrographs of *via1* leaves showcase a prominently reduced thylakoid network and accumulation of vesicular structure. A corresponding area from the wild-type plant is provided for comparison. Squares in each image represent the area that was acquired with higher magnifications. **(B)** 3D reconstruction of chloroplasts using serial block face scanning electron microscopy. Young leaf tissue from both wild-type and *via1* plants was collected after 72 h of continuous light exposure and imaged using a FEI Apreo VolumeScope from Thermo Fisher Scientific. The outer envelope of the chloroplasts was manually segmented and is represented in green, highlighting the diverse chloroplast morphologies observed between the two genotypes. Complete segmented stacks are available in the Suppl. Videos S1 and S2.

## Discussion

ESCRT-mediated membrane remodeling is a conserved phenomenon in the cytoplasm of all eukaryotic cells. However, the recent biochemical characterization of eukaryotic-like ESCRT-I, ESCRT-II, and ESCRT-III components in Asgard archaea suggests a pre-eukaryotic origin for the ESCRT machinery (42–44). In line with this hypothesis, recent structural analyses have highlighted PspA, a bacterial protein important for membrane integrity, and VIPP1, a critical protein for thylakoid biogenesis in cyanobacteria and chloroplasts of photosynthetic eukaryotes, as members of the ESCRT-III family of proteins, capable of inducing membrane deformation upon binding (18, 19, 21).

In this work, we provide further evidence supporting the ancient evolutionary origin of the ESCRT system by showing similarities between ESCRT-II proteins and VIA1, the chloroplast and cyanobacterial membrane protein we described here. We found that VIA1 interacts with VIPP1 in two distantly related photosynthetic eukaryotes, the unicellular green alga *C. reinhardtii* and the flowering plant *A. thaliana* (Figure 1, 4). Structural prediction analyses indicate the presence of three transmembrane helices within VIA1, accompanied by a WH domain, a distinctive feature of ESCRT-II proteins (Figure 5). Our analyses suggest that VIA1’s WH domain may interact with VIPP1’s first alpha helix, consistent with an evolutionary conservation of the typical interface for ESCRT-II-ESCRT-III interactions (39).

We also provide insights regarding the functional importance of the ancient ESCRT-II-like proteins *in vivo*—a challenging task in archaea due to the absence of established genetic tools and the slow growth rate of these organisms in standard laboratory conditions (45). In particular, we show that deletion of VIA1 leads to drastic changes in thylakoid membrane morphology in both Chlamydomonas and Arabidopsis. Specifically, the absence of VIA1 in Arabidopsis results in a severe but transient reduction of the thylakoid membrane network in chloroplasts of newly formed leaf tissue (Figure 9A). The chloroplasts in this developing tissue were misshaped and contained numerous vesicular structures and membrane invaginations (Figure 9B). In contrast, the effects of *VIA1* deletion in Chlamydomonas manifest primarily under high-light stress conditions, where thylakoids underwent swelling before any defect in the photosynthetic machinery became apparent (Figure 6E, 7).

Based on these observations and previous reports regarding the function of VIPP1, we speculate that VIA1 functions together with VIPP1 in the formation of lipid vesicles and invaginations originating from the inner envelope of chloroplasts in Arabidopsis. This hypothesis could explain why VIA1’s function is necessary during early leaf development, possibly due to high demand for the machinery required for chloroplast division, and consequently thylakoid biogenesis, in these rapidly expanding cells. In the case of Chlamydomonas, the role of VIA1 in thylakoid membrane biogenesis is less clear, because thylakoids in mutants lacking this protein are indistinguishable from those in WT under standard light conditions. However, when exposed to prolonged high light, the lack of VIA1 has a clear impact on chloroplast membrane morphology, causing thylakoid swelling. In turn, this defect makes the cells more susceptible to such stress. Thus, in this unicellular alga, VIA1 appears to primarily preserve thylakoid membrane integrity during photooxidative stress, possibly by affecting VIPP1 activity. In this regard, VIA1 resembles LEM2, a two-pass membrane protein located in the inner nuclear envelope that recruits the ESCRT-II/ESCRT-III hybrid protein CHMP7 and other ESCRT-III proteins to seal holes in reforming nuclear membranes (46). The notion that VIA1 may be involved in the homeostasis and maintenance of damaged chloroplast membranes under photooxidative stress is also supported by the observation of a similar thylakoid swelling phenotype in Chlamydomonas cells following VIPP1 knockdown under excessive light conditions (14). Furthermore, this idea provides a plausible explanation for the strong transcriptional induction of VIA1 observed in response to high light exposure (47).

Since the eukaryotic ESCRT system comprises several multi-protein complexes, it is also conceivable that other molecular factors are involved in thylakoid membrane biogenesis and maintenance. Some of these proteins might belong to the list of conserved VIPP1-interacting partners we identified. This list includes the Chlamydomonas protein CPLD42, whose Arabidopsis counterpart FLAP1 has been proposed to play a role in acclimation to fluctuating light conditions (31), a predicted PPR superfamily protein, and a small transmembrane domain-containing protein of unknown function. These proteins remained undetected in a recent VIPP1 proximity-labeling study conducted in Chlamydomonas (28). Thus, it appears that AP-MS and proximity-labeling methods can be complementary in identifying the components of multi-protein complexes. Combined together, they offer a comprehensive overview of the VIPP1-interactome in chloroplasts of alga and plant cells.

Overall, our findings reveal the involvement of an ESCRT-II-like protein, VIA1, in thylakoid membrane biogenesis and maintenance. This, alongside the identification of other evolutionarily conserved proteins that interact with the ESCRT-III-like VIPP1 protein, advances our understanding of the evolutionary history, protein composition, and molecular mechanism of the thylakoid membrane remodeling system.

## Materials and Methods

### Plant Growth conditions

All plants were grown on soil in a controlled environment chamber (Percival AR-95L3; CLF Plant Climatics) fitted with fluorescent lamps (Philips 368290 F25T8/TL841 ALTO) and supplemented with red LED panels, programmed for a 12-h light / 12-h dark cycle, unless otherwise specified. The light intensity was 150 μmol photons m^−2^s^−1^. The temperature was 20°C with 60% relative humidity. For experiments on plates, seedlings were grown on 1.2% [w/v] agar plates containing half-strength Murashige and Skoog medium (Duchefa Biochemie) and incubated in a walk-in chamber (Kälte 3000 AG) with similar light and temperature settings as above.

### Molecular Cloning

The attB1- and attB2-flanked coding sequence of *At*VIPP1 were amplified using Arabidopsis cDNA as a template and inserted into the gateway-compatible entry vector pDONR221 using the BP Clonase II enzyme mix (Invitrogen). The resulting pDONR221-*At*VIPP1 entry vectors were then recombined into the gateway-compatible destination vectors, pB7YWG2 or pH7CWG2 (48, 49), with a constitutive CaMV 35S promoter and either a YFP or CFP C-terminal tag using the LR Clonase II enzyme mix (Invitrogen). The YFP-tagged construct was used for the generation of stable transgenic Arabidopsis lines of and co-IP experiments, while the CFP-tagged version was used for sub-cellular localization assays.

Similarly, the coding sequence of *At*VIA1 was amplified, cloned in the pDONR221 Gateway entry vector, and further recombined with the destination vector pUBC-YFP-dest (50) with a constitutive UBQ10 promoter and a C-terminal YFP tag. This entry vector was additionally recombined with the destination expression vector pH7m34GW, together with the UBQ10 promoter and a C-terminal 3xFLAG tag. Both constructs were used for co-IP assays in *N. benthamiana,* while the YFP-tagged *At*VIA1 construct was also used for localization assays. For the generation of *via1/pro*VIA1:VIA1FLAG lines, an attB1- and attB2-flanked genomic fragment of *At*VIA1, including the promoter and 5′ UTR (648 bp upstream of the start codon), was amplified from Arabidopsis genomic DNA and cloned into the gateway-compatible pDONR221 vector via BP reaction. This entry vector was recombined into the destination vector pH7m34GW together with the 3xFLAG tag at the C-terminus using Gateway LR reaction. All expression constructs were transformed into *Agrobacterium tumefaciens* strain GV3101 (pMP90) (51) via electroporation and subsequently used for infiltration of *Nicotiana benthamiana*.

### Arabidopsis transformation

For stable expression of proteins in Arabidopsis, the floral-dip method of transformation was utilized as described by (52). Harvested seeds were surface sterilized with 70% [v/v] ethanol and germinated on soil or agar plates, and transformants were selected based on resistance to Basta® herbicide (sprayed 0.1% [v/v], Sigma Aldrich) or on ½ strength MS-plates containing 25 µg/mL Hygromycin B (Carl Roth), depending on the construct.

### Immunoprecipitation and mass spectrometry (*A. thaliana*)

Total protein was extracted from leaves harvested from 35-day-old WT Col-0 Arabidopsis plants constitutively expressing FNR-GFP (Marques et al., 2004) or VIPP1-YFP by homogenizing the leaves in extraction buffer (50 mM Tris-HCl pH 8.0, 150 mM NaCl, 1 mM Dithiothreitol (DTT) 1% [v/v] Triton X-100, and 1X Complete Protease Inhibitor Cocktail, Roche) at a ratio of 1 mL buffer per 250 mg fresh weight using a Dounce homogenizer. Cellular debris was removed by centrifugation twice at 19,000 g for 5 min at 4°C. The supernatant was incubated with 25 μL of pre-equilibrated ChromoTek GFP-Trap magnetic agarose beads (ChromoTek) for 1 h, at 4°C on a rotating wheel. After incubation, the beads were retrieved using a magnetic separation rack and washed twice with wash buffer (50 mM Tris-HCl pH 8.0, 150 mM NaCl, 1 mM DTT, 0.5% [v/v] NP-40, and 1X Complete Protease Inhibitor Cocktail (Roche)) and once with dilution buffer (50 mM Tris-HCl pH 8.0, 150 mM NaCl). Beads were further washed 3 times with digest buffer (10 mM Tris, pH 8.2,2 mM CaCl_2_). For reduction and alkylation, the beads were covered in lysis buffer, and Tris(2-carboxyethyl)phosphine and 2-Chloroacetamide was added to a final concentration of 2 mM and 15 mM, respectively, and incubated for 30 min at 30°C at 700 rpm. Trypsin was added to enzymatically digest peptides, assisted with Microwave at 60°C for 30 min. Supernatants were collected, and the remining peptides were extracted from the beads with 0.1% [v/v] trifluoroacetic acid and 50% [v/v] acetonitrile. The supernatants were dried in speed vac and then dissolved in aqueous 3% [v/v] acetonitrile with 0.1% [v/v] formic acid. Peptide concentration was estimated with a Lunatic UV/Vis absorbance spectrometer (Unchained Lab). Peptides were separated on an M-class UPLC (Waters^TM^) running in single-pump trapping 75-um scale configuration incl. a nanoEase M/Z HSS T3 Column (analytical column of 100Å pore size, 1.8 µm particle size, inner diameter and length:75 µm х 250 mm) and analyzed on an Orbitrap Exploris 480 mass spectrometer (Thermo Scientific) operated on DDA mode.

The LC-MS data was processed using using the Philosopher toolkit including MSFragger as a search engine followed by IonQuant (53–55). Protein quantification results were extracted from the combined_protein.tsv file. We used the columns with the intensity suffix, which stores the normalized protein intensity using the sequences after razor assignment. SaintExpress software (56) was used to score potential interactions between observed proteins (potential prey) and the bait protein. Scaffold 5 (Proteome Software) was used to validate MS/MS-based peptide identifications and score the semi-quantitative value.

### Co-immunoprecipitation and immunoblotting (*N. benthamiana* leaf tissue)

Total protein was extracted from 4-week-old *N. benthamiana* leaves transiently expressing *At*VIPP1-FLAG, *At*VIA1-YFP, *At*VIA1-FLAG, or *Cp*FeS-YFP under control of the Ubiquitin10 (UBQ10pro) or CaMV 35S promoter. Immunoprecipitation was performed as described previously (57). For immunoblotting, a mix of anti-GFP and anti-FLAG primary antibodies were used. Subsequently, anti-rabbit IRDye 800CW and anti-mouse IRDye 680CW secondary antibodies (1:10,000, polyclonal/donkey, LI-COR Biosciences) were used for detection of signals specific to antigen-antibody binding with an Odyssey CLx imaging system (LI-COR Biosciences) as described previously (58).

### Confocal laser scanning microscopy (CLSM)

Agrobacterium-mediated transient transformation of *N. benthamiana* leaf tissue was performed as described previously (59). The transiently transformed leaf cells were analyzed using a confocal system, LSM780 from Zeiss using either laser of 514 nm (YFP), 458 nm (CFP) and 633 nm (chlorophyll) wavelengths. Image acquisition was done sequentially for different channels using filters ranging from 526 to 624 nm (YFP), 463 to 509 nm (CFP), and 647 to 721 nm (chlorophyll). Images were acquired in several Z-stacks, and projected for their maximum intensity for visualization.

### Phylogenetic analyses

For phylogenetic analysis, amino acid sequences of *At*VIA1 homologs were retrieved from NCBI by querying AtVIA1 using the blastp suite of NCBI-BLAST against the refseq_protein and reseq_select databases. Sequences were manually selected so that the major evolutionary branches were represented. Evolutionary analyses were conducted in MEGA7 (60) as described by Hall (61). In total, VIA1 sequences from 140 species were aligned using the MUSCLE algorithm (62). The best substitution model was determined using a feature in MEGA7 and the LG+G+F model was chosen due to the highest BIC score. With these parameters, a Maximum Likelihood tree was constructed with complete elimination of gaps and missing data. A discrete Gamma distribution was used to model evolutionary rate differences among sites (5 categories (+G, parameter = 1.2033)). To estimate the reliability of the constructed tree, 1000 bootstrap replications were performed and included to represent the evolutionary history of VIA1. Branches corresponding to partitions reproduced in less than 50% bootstrap replicates are collapsed. For visualization of the phylogenetic tree, the iTOL web-based tool was utilized (63).

### Growth conditions (*C. reinhardtii*)

*Chlamydomonas reinhardtii* strains were maintained on solid Tris-Acetate-Phosphate (TAP) media containing 1.6% [w/v] agar (USP grade, Thermo Fisher Scientific) and Hutner’s trace elements (64) (Chlamydomonas Resource Center) at 22°C under low light conditions (∼30 µmol photons m^−2^ s^−1^). Mutant strains harboring the mutagenic cassette (CrRS432, CrRS433) were maintained on solid TAP media supplemented with 10 µg mL^−1^ paromomycin. Unless otherwise stated, strains were grown in liquid culture under standard light conditions with an intensity of 100 µmol photons m⁻² s⁻¹ and in the absence of any antibiotics. A list of all strains used in this research can be found in Table S3.

### Co-Immunoprecipitation and Mass spectrometry (*C. reinhardtii*)

A 5 mL volume of Tris-Acetate-Phosphate (TAP) medium was initially inoculated with wild-type (WT) cells and cultivated until the cell density reached 2-4 x 10⁶ cells mL⁻¹. Subsequently, the culture was progressively expanded through sequential dilution to 50 mL and 150 mL volumes. Eventually, the culture was transferred to 2-L flasks and further diluted by introducing 750 mL of TAP medium. After a 24-h incubation, the cells were harvested by centrifugation and the resulting pellet resuspended in a 1:1 ratio with ice-cold 2XIP buffer (100 mM HEPES, 100 mM Potassium Acetate, 4 mM Magnesium Acetate, 2 mM Calcium Chloride, and 400 mM sorbitol) freshly supplemented with Protease Inhibitors EDTA-free (Roche). The resuspended mixture was added dropwise into a 50 mL falcon containing liquid nitrogen in a liquid nitrogen bath. The resulting 5-mm Chlamydomonas pellets were homogenized in a pre-chilled cryo-mill (Retsch) via 5 cycles of 3-min milling at a frequency of 20 s⁻¹, alternating with 2-min pauses to re-cool the vials by immersion in liquid nitrogen. The cell powder was transferred into a ice-cold 50 mL Falcon tube to thaw. The cell lysate was further processed by douncing it 30 times using a Kontes Dual 21 homogenizer. Subsequently, 500 µL of lysate was combined with 500 µl of cold IP buffer and 110 µl of 10% [w/v] digitonin. The mixture was incubated on a rotating wheel (1 h, 4°C). The solubilized lysate was clarified through centrifugation at 21,300 x g for 30 minutes at 4°C.

For immunoprecipitation (IP), 150 µL of protein A Dynabeads were used per sample. The beads were thoroughly washed 4 times with 1 mL of IP buffer, then resuspended in their original volume and 30 µl of 1 mg mL^-1^ VIPP1 antibody per sample was added. Additional samples were prepared by incubating lysates with non-coupled beads, as negative controls. The beads were placed on a rotating wheel for 45 min at 20°C, transferred to a magnetic stand (MagnaRack) to remove the supernatant, then washed once with 1 mL of IP buffer, and twice with 1 mL of IP buffer + 0.1% [w/v] digitonin (for 5 min at 4°C). At this point, 750 µl of clarified lysate was added to the antibody-bound beads, and incubated on a rotating wheel (3 h, 4°C). After placing the beads on the magnetic stand, the flowthrough was removed, and 4 washes with 1 mL of IP buffer + 0.1% [w/v] digitonin were performed (the first two lasting 5 min each and the last two lasting 10 min each, all at 4°C). For the elution step, denaturing conditions were employed. Specifically, the beads were resuspended in 60 µL of 2X denaturing lysis buffer (100 mM Tris-HCl pH 8.0, 600 mM NaCl, 4% [w/v] SDS, 20 mM EDTA) and incubated at 70°C for 15 min in a thermomixer at 850 rpm. Once placed on the magnetic stand, the supernatant was separated and stored at −20°C. For mass spectrometry analysis, the eluates underwent further preparation. Firstly, 5 µl of 5X SDS-loading buffer, composed of 250 mM Tris-HCl pH 6.8, 5% [w/v] SDS, 0.025% [w/v] bromophenol blue, and 25% [v/v] glycerol, freshly supplemented with 5% [v/v] 2-mercaptoethanol, was added to 20 µl of the denatured eluates. The resulting mixtures were then incubated for 5 min at 95°C. Subsequently, the proteins samples were separated by SDS-PAGE and stained with colloidal Coomassie solution. Sample lanes were excised from the gel, transferred into 1.5 mL tubes containing 1% acetic acid and stored at 4°C until use.

### Sample processing for Mass Spectrometry (MS) (*C. reinhardtii*)

Coomassie-stained gel bands were cut into 2-3 mm pieces, transferred to 0.6-mL tubes, and incubated with different solutions by shaking for 10 min at 20°C followed by removal of the supernatant as follows: Gel pieces were washed with 200 µl 100 mM ammonium bicarbonate (ABC), destained by 2 repeated rounds of shrinking in 200 µl 50% acetonitrile in 50 mM ABC and reswelling in 200 µl 100 mM ABC. Gel pieces were shrunk with 100 µL acetonitrile before being reduced with 100 µl of 6 mM DTT in 100 mM ABC by incubation at 57°C for 30 min and alkylated with 100 µl of 28 mM MMTS in 100 mM ABC by incubation at RT for 30 min.

Wash steps were repeated as described for de-staining and gel pieces were shortly dried in a speed-vac after the final shrinking step. Gel pieces were soaked in 12.5 ng/µl Trypsin in ABC for 5 min at 4°C. Excess solution was removed, ABC was added to cover the pieces, and samples were kept overnight at 37°C. The supernatant containing tryptic peptides was transferred to a fresh tube and gel pieces were extracted by addition of 20 µL 5% [v/v] formic acid and sonication for 10 min in a cooled ultrasonic bath. This step was performed twice. All supernatants were then combined. A similar aliquot of each digest was analyzed by LC-MS/MS.

### NanoLC-MS/MS Analysis (*C. reinhardtii*)

The nano HPLC system (UltiMate 3000 RSLC nano system or Vanquish Neo UHPLC, Thermo Fisher Scientific) was coupled to an Exploris 480 mass spectrometer equipped with a FAIMS pro interface and a Nanospray Flex ion source (Thermo Fisher Scientific). Peptides were loaded onto a trap column (PepMap Acclaim C18, 5 mm × 300 μm ID, 5 μm particles, 100 Å pore size, Thermo Fisher Scientific) using 0.1% [v/v] trifluoroacetic acid as mobile phase. After loading, the trap column was switched in line with the analytical column (PepMap Acclaim C18, 500 mm × 75 μm ID, 2 μm, 100 Å, Thermo Fisher Scientific). Peptides were eluted using a flow rate of 230 nl/min, starting with the mobile phases 98% [v/v] A (0.1% [v/v] formic acid in water) and 2% [v/v] B (80% [v/v] acetonitrile, 0.1% [v/v] formic acid) and linearly increasing to 35% [v/v] B over the next 120 min. This was followed by a steep gradient to 90% [v/v] B over 5 min, held for 5 min, and ramped down in 2 min to the starting conditions of 98% [v/v] A and 2% [v/v] B for equilibration at 30°C.

The mass spectrometer was operated in data-dependent mode, performing a full scan (m/z range 350-1200, resolution 60,000, normalized AGC target 1E6) at 3 different compensation voltages (CV-45, −60, −75), each followed by MS/MS scans of the most abundant ions for a cycle time of 0.9 second for CV −45 and −60 and 0.7 sec for CV-75. MS/MS spectra were acquired using a collision energy of 30, isolation width of 1.0 m/z, resolution of 30.000, max fill time 100 ms, normalized AGC target of 2E5 and intensity threshold of 2.5E4. Precursor ions selected for fragmentation (include charge state 2-6) were excluded for 45 s.

### MS Data Processing (*C. reinhardtii*)

For peptide identification, the RAW-files were loaded into Proteome Discoverer (version 2.5.0.400, Thermo Scientific). All MS/MS spectra were searched using MSAmanda v2.0.0.19924 (65). The peptide and fragment mass tolerance were set to ±10 ppm, the maximum number of missed cleavages was set to 2, using tryptic enzymatic specificity without proline restriction. Peptide and protein identification was performed in two steps.

For an initial search, the RAW-files were searched against the database Chlamydomonas_reinhardtii_v6_Phytozome.fasta (30,378 sequences; 135,270,025 residues), supplemented with common contaminants and sequences of tagged proteins of interest using Iodoacetamide derivative on cysteine as a fixed modification. The result was filtered to 1 % FDR on protein level using the Percolator algorithm (66) integrated in Proteome Discoverer. A sub-database of proteins identified in this search was generated for further processing. For the second search, the RAW-files were searched against the created sub-database using the same settings as above and considering the following additional variable modifications: oxidation on methionine, phosphorylation on serine, threonine and tyrosine, deamidation on asparagine and glutamine, ubiquitination on lysine, and glutamine to pyro-glutamate conversion at peptide N-terminal glutamine, acetylation on protein N-terminus. The localization of the post-translational modification sites within the peptides was performed with the tool ptmRS, based on the tool phosphoRS (67).

Identifications were filtered again to 1% FDR at the protein and PSM level and, additionally, an Amanda score cut-off of at least 150 was applied. Proteins were filtered to be identified by a minimum of 2 PSMs in at least 1 sample and quantified based on at least 3 quantified peptides over all samples. Peptides were subjected to label-free quantification using IMP-apQuant (68). Proteins were quantified by summing unique and razor peptides and applying intensity-based absolute quantification (69). Statistical significance of differentially expressed proteins was determined using limma (70).

### VIA1 structure modelling and alignment

For the structural analysis of *At*VIA1 and *Cr*VIA1, the transit peptide sequences were predicted using TargetP 2.0 (71), and transmembrane domains using InterProScan (72) and DeepTMHMM (73). The template search for the structural modeling of *At*VIA1 was performed via SWISS-MODEL (74) and Phyre2 (37), using the full-length amino acid sequence of *At*VIA1 without the transit peptide. In addition, the predicted tertiary protein structure of *At*VIA1 and *Cr*VIA1 were modeled by AlphaFold2 (75) and the resulting structural model was used to search for related structures using the BackPhyre function of Phyre2 (37). Structural prediction of protein complexes was carried out via AlphaFold2 using the multimer_v3 model type. Graphical modifications on the protein structures were done using PyMOL v2.5 (76) and ChimeraX-1.4 (77). The structural alignment between AtVIA1 and CrVIA1 was carried out using Pairwise Structure Alignment, an online tool from RCSB-PDB (36).

### Isolation and genotyping of *C. reinhardtii VIA1* insertion lines

Genomic DNA extraction from WT, *via1-1* and *via1-2* liquid cultures in exponential growth phase and in standard light conditions (∼80 µmol photons m^−2^ s^−1^) was carried out as previously described (47). The insertion of the mutagenic cassette in the *CrVIA1* locus was verified by PCR in each mutant. All PCRs were performed using KOD Hot Start DNA Polymerase (EMD Millipore). In the *via-1* strain, the insertion occurs in exon 10. Therefore, primer pairs oRS39 and oRS40 were designed to anneal in exon 9 and the 3’ untranslated region, respectively (see Fig. S3 for detail). This primer pair allows the amplification of the WT locus as well as of the mutated locus containing the cassette (PCR conditions: 95°C 2 min, 95°C 20 sec, 55°C 10 sec, 70°C 1 min 40 sec). To selectively amplify a section of the cassette inserted in the *CrVIA1* locus, the oRS01 primer was designed to anneal within the mutagenic cassette and was used in conjunction with oRS39 (PCR conditions: 95°C 2 min, 95°C 20 sec, 55°C 10 sec, 70°C 28 sec).

For *via-2*, which harbors the insertion within intron 7, primer pairs oRS37 and oRS38 were designed to anneal in exon 6 and exon 8, respectively. This primer pair allows the amplification of the WT locus as well as of the mutated locus containing the mutagenic cassette (PCR conditions: 95°C 2 min, 95°C 20 sec, 55°C 10 sec, 70°C 1 min 40 sec). Moreover, to selectively amplify a section of the cassette inserted in the *CrVIA1* locus, the oRS30 primer was designed to anneal within the mutagenic cassette and was used in conjunction with oRS38 (PCR conditions: 95°C 2 min, 95°C 20 sec, 55°C 10 sec, 70°C 30 sec). To ensure the quality of the employed genomic DNA (gDNA), a mating type-specific PCR (78) was performed as a control. All PCR reactions were subjected to electrophoresis on 1% [w/v] agarose gels to verify their expected sizes.

### Backcross and tetrad dissection analysis

Cells were streaked onto a fresh TAP agar plate and grown under light (100 µmol photons m^-2^ s^-1^) for approximately 3-4 days. The cells were subsequently restreaked onto a TAP agar plate containing 1/10 of the standard NH_4_Cl concentration to induce starvation. The day after, gametes were suspended in 100 µL of H_2_O, achieving a concentration of 10^6^ cells mL^-1^, and agitated for 30-60 minutes under light (100 µmol photons m^-2^ s^-1^). Mating efficiency was monitored every 30 minutes by mixing 2 µL of the mating mixture with 8 µL of Lugol’s solution (Gatt-Koller) and observing quadriflagellate formation under a microscope. Generally, mating occurred within the second hour. Subsequently, 100 µL of the mating mixture was spotted in the middle of a TAP 4% [w/v] agar plate, which was covered with aluminum foil. After six days, vegetative cells were gently scraped off using a rectangular razor blade (Apollo Herkenrath), while the zygotes remained adhered to the agar surface. 100 µL of liquid TAP was applied over the zygotes and a razor blade (Aesculap AG) was used to detach them. The resulting mixture was transferred to a fresh TAP 1.6% [w/v] agar plate, drawing a line onto the center of it. The plate was then incubated under light (100 µmol photons m^-2^ s^-1^) for 24 h to allow germination. A dissection microscope (SporePlay+) was employed to identify and dissect tetrads. Once visible to the naked eye, the tetrads were re-arrayed into a rectangular agar plate (Singer Instruments) in a 96-array format and replicated using TAP plates or TAP + paromomycin plates.

### RNA extraction and qRT-PCR for gene expression analysis (*C. reinhardtii*)

RNA extraction from WT, *via1-1*, and *via1-2* liquid cultures in exponential growth phase and in standard light conditions (∼80 µmol photons m^−2^ s^−1^) was carried out as described (47). Total RNA (1.65 µg) and oligo(dT)_18_ were used to synthesize cDNA using a cDNA synthesis kit (Maxima First Strand cDNA, Thermo Fisher Scientific), following the manufacturer’s instructions. cDNA was diluted ten-fold prior to qPCR analysis, which was carried out employing SYBR green (BioRad) as per the manufacturer’s instructions. Primers to amplify the *VIA1* transcript are listed in Table S1. To control for DNA contamination, primer dimers, or misannealing, the same volume of the master mix without cDNA was added to a well of the 96-well plate. Raw Ct values were analyzed per the ‘eleven golden rules’, as described in Udvardi et al., 2008. *GBLP* (Cre06.g278222) was chosen as a reference housekeeping transcript during normalization. One biological replicate was analyzed for *via-1*, and two biological replicates were analyzed for *via2-2*. Additionally, each experiment included two technical replicates.

### Chlorophyll Fluorescence Measurements (*C. reinhardtii*)

Chlorophyll fluorescence measurements of Chlamydomonas were performed using a Dual PAM 100 fluorometer (Walz, Germany). Cells were grown in minimum media for 5 days and harvested during the exponential growth phase and subsequently subjected to a 30-minute period of dark adaptation while maintaining gentle agitation. The following parameters were calculated: Photosystem I Quantum Yield ((Pm - Pm’)/Pm), Photosystem II Quantum Yield (Fv/Fm) and Non-Photochemical Quenching ((Fm - Fm’)/Fm’) where Fm’ is the maximal chlorophyll fluorescence yield during exposure to light. Calculations were based on established methodologies for measurement of photosynthetic parameters (79, 80).

### High-light assay and time course (*C. reinhardtii*)

Liquid cultures were initiated from freshly streaked colonies (3-4 days old) on TAP plates kept under continuous light (100 µmol photons m^-2^ s^-1^). A loop of cells was resuspended in 15 mL of TAP media using 50 mL glass flask. Cells were grown for 2-3 days under continuous light (100 µmol photons m^-2^ s^-1^) on a shaker set at 130 rpm. Upon reaching a cell density of 10^6^ cells mL^-1^, the cultures were scaled up to 50 mL and allowed to grow for an additional 1-2 days. Subsequently, the total chlorophyll concentration of each culture was measured following the methanol extraction method described previously (81). Cells were equally diluted to 8 µg chlorophyll mL^−1^ in a final volume of 31 mL in 50 mL Falcon tubes. Three replicates were generated for each strain. The high-light (HL) treatment was executed with an illuminance of ∼800 µmol photons m^-2^ s^-1^, employing a Plant LED Grow Light (Phlizon, Cob Series 3000W) placed approximately 40 cm away from the cultures. To ensure uniform photon exposure across all samples, light intensity was measured along each tube. Additionally, to prevent bias, cultures were randomly rotated every 24 h during the treatment. A temperature increase of 4-5°C was observed upon the transfer of the cultures to HL conditions. During the high light assay, images and chlorophyll measurements were taken daily from the beginning of the experiment, for a total of three days. For the high-light time course immunoblot analysis, the HL treatment was executed in the same way for a duration of 8 h. In this case, at the onset of the HL treatment, cells were diluted in a final volume of 100 mL and 5 mL aliquots were collected at 2-h intervals throughout the course of the treatment. Cells were centrifuged at 3,000 x g for 5 minutes, the supernatant was discarded, and the resultant cell pellet was stored at −20°C for total protein extraction.

### Total protein extraction and immunoblot analysis (*C. reinhardtii*)

For all experiments, denaturing protein extraction from whole-cell lysate was conducted via a denaturing SDS extraction procedure. Cells from a 5 mL culture in the exponential growth phase were collected at 3,000 x g for 5 minutes and resuspended in 150 µl of SDS-lysis buffer (100 mM Tris-HCl pH 8.0, 600 mM NaCl, 4% [w/v] SDS, 20 mM EDTA) freshly supplemented with Protease Inhibitors EDTA-free (Roche). The samples were vortexed for 10 minutes at room temperature (RT), incubated for 30 minutes at 37 °C, and then centrifuged at 21,300 x g for 15 minutes at RT to eliminate cellular debris. The supernatant was then transferred to a new Eppendorf tube and supplemented with ¼ volume of 5X SDS-loading buffer (250 mM Tris-HCl pH 6.8, 5% [w/v] SDS, 0.025% [w/v] bromophenol blue, and 25% [v/v] glycerol) freshly supplemented with 5% [v/v] 2-mercaptoethanol. A 5 µL aliquot from the supernatant was kept for protein quantification, which was performed using a BCA assay. Specifically, 2 µl of the protein extract was mixed with 200 µl of BCA/copper sulfate solution (at a 1:50 ratio). After incubation for 5 minutes at 50°C, the absorbance at 562 nm was measured to estimate the protein concentration, relying on a BSA standard curve.

For subsequent immunoblot analysis, 40 µg of the denatured protein extract was employed. Proteins were separated via SDS-PAGE utilizing Criterion Precast Gels (Bio-Rad) and were transferred onto a nitrocellulose membrane, pore size 0.2 μm (Amersham). A PBS-T solution supplemented with 5% [w/v] instant nonfat dry milk (Maresi) was employed for blocking non-specific signals. This blocking step was carried out for 1 h at RT. Both primary and secondary antibodies were diluted in this blocking buffer (Table S2). The incubation step with the primary antibody step was conducted for 1 h at RT, except in the cases of anti-D1 where it was extended overnight at 4°C.

To detect primary antibodies, HRP-conjugated anti-rabbit or anti-mouse secondary antibodies were employed at a dilution of 1:10,000 at RT for 1 h. Following each incubation step, membranes were washed three times for 10 minutes at RT using PBS-T supplemented with 1% [w/v] instant nonfat dry milk. Finally, membranes were rinsed with water, and the signal was developed using a luminol-based enhanced chemiluminescence (ECL) method (SuperSignal West Dura or SuperSignal West Femto, ThermoFisher Scientific).

### Transmission electron microscopy (*C. reinhardtii*)

Wild-type (WT) and *via1-2* cell cultures were subjected to HL for 0, 2, 4, and 6 h. 10 mL aliquots from each time point, at a concentration of approximately 10^6^ cells mL^-1^, were gently spun down at 500 g for 3 minutes, and the resulting cell pellets were fixed using a mixture of 2% [v/v] paraformaldehyde (Electron Microscopy Sciences, Hatfield, PA) and 2% [v/v] glutaraldehyde (Agar Scientific, Essex, UK) in 0.1 M sodium cacodylate buffer, pH 7.4. This fixation process was carried out overnight at room temperature. Samples were then rinsed with the same buffer, post-fixed in 2% [w/v] osmium tetroxide (Agar Scientific, Essex, UK) in 0.1 M sodium cacodylate buffer, pH 7.4, dehydrated in a graded series of acetone and embedded in Agar 100 resin (Agar Scientific, Essex, UK). Subsequently, 70 nm sections were cut and post-stained with 2% uranyl acetate [w/v] and Reynolds lead citrate (Delta Microscopies, Mauressac, France). Micrographs were recorded on a FEI Morgagni 268D (FEI, Eindhoven, The Netherlands) operated at 80 keV, equipped with a Mega View III CCD camera (Olympus-SIS).

### Isolation and genotyping of A. *thaliana* T-DNA insertion lines

The T-DNA insertion line *via1-1* (SALK_057879C) was obtained from the Nottingham Arabidopsis Stock Centre (NASC). Segregating populations were screened by PCR-based genotyping to select homozygous plants for the T-DNA insertion. Genomic DNA was isolated as described (82). Two genomic primers, VIA1_LP and VIA1_RP, were utilized to amplify the genomic region while the VIA1_RP primer and one primer matching the T-DNA left border sequence (SALK_Lba1) were used to amplify the genomic region between the gene of interest and T-DNA insert using DreamTaq DNA Polymerase (Thermo Fischer Scientific) at a melting temperature of 55°C, as per the manufacturer’s instructions.

### RNA Extraction and RT-PCR (*A. thaliana*)

For RNA extraction, 3-week-old Arabidopsis rosettes grown on ½ strength MS plates were snap-frozen in liquid nitrogen and homogenized using glass beads. Two rosettes were pooled together to achieve a fresh weight of 10 mg per sample. Total RNA was isolated using the GENEzol extraction buffer according to the manufacturer’s guidelines (Geneaid Biotech). Contamination of genomic DNA was removed by treating the samples with DNase I (Roche). First-strand cDNAs were synthesized from 2 µg of RNA using oligo(dT)_18_ primers and the RevertAid First Strand cDNA Synthesis Kit (Thermo Fisher Scientific). The resulting cDNA was used for RT-PCR reactions performed with control primers for the housekeeping gene ACTIN and gene-specific primers for VIA1. Primers for RT-PCR were designed using AtRTPrimer web-tool (83) and sequences are listed in the supplemental dataset.

### Photosynthetic measurement (*A. thaliana*)

The maximum quantum yield of Photosystem II was measured as a function of the ratio of variable to maximum fluorescence (Fv/Fm). Each plant was dark-adapted for at least 5 minutes directly before the measurement. Chlorophyll fluorescence measurements were conducted using a FluorCam 800MF equipped with FluorCam 7 software from Photon Systems Instruments. The software’s default program “Fv/Fm” was used, with actinic light intensity set at 20% and saturation light intensity at 100%. For 28-day-old plants, at least three biological replicates were imaged, and the experiment was repeated three times. Similarly, imaging of seedlings was repeated twice, with at least 15 individual 10-14-day-old Arabidopsis seedlings grown on ½ strength MS plates in each case. Statistical significance testing was performed using Welch’s t-test in GraphPad Prism software to assess variations.

### Sample preparation for TEM imaging (*A. thaliana*)

Leaf sections were harvested in the morning after receiving 90 minutes of light or 96 h of continuous light. Samples were chemically fixed in fixation buffer (2.5% [v/v] glutaraldehyde, 2% [w/v] formaldehyde in 0.1 M sodium cacodylate pH 7.4), washed three times with 0.1 M sodium cacodylate pH 7.4, stained with 2% [w/v] osmium tetroxide, and embedded in epoxy resin (Spurr Low Viscosity Embedding Kit, Polysciences Inc) as described earlier (84). Eventually, the samples were cured in silicone molds for at least 24 h at 60°C. All steps prior to curing were microwave-assisted using a BioWave Pro+ (Ted Pella), with 40 s/250 W settings for dehydration steps and 5 min/200 W/T_max_ = 22°C settings for embedding. The resulting resin blocks were sectioned using an ultramicrotome, and ultra-thin sections (50 nm) were transferred onto formvar-coated copper grids (EMS). These sections were stained with uranyl acetate for 10 minutes, washed three times with water, and stained with lead citrate for another 10 minutes, and then washed three times with water. Samples were subsequently imaged with a transmission electron microscope TFS Morgagni 268 (FEI Company) at 100 kV. Shading correction in the acquired images was done using thresholding tool of Fiji (85).

### SBF-SEM (Serial Block Face-Scanning Electron Microscopy)

Sections from newly emerging leaf base of 10-day-old Arabidopsis seedlings, grown under continuous light for an additional 72 h, were harvested, chemically fixed in 2.5% [v/v] paraformaldehyde and 2% [v/v] Glutaraldehyde in 0.1 M sodium cacodylate buffer, pH 7.4, stained with 2% [w/v] osmium tetroxide, 2% [w/v] uranyl acetate and lead citrate and embedded into Durcupan™ ACM epoxy resin (Sigma-Aldrich) as described (86). The resultant blocks were trimmed, glued on stubs, coated with gold/palladium, and sectioned/scanned approximately 500 times using a scanning electron microscope (FEI Apreo VolumeScope, Thermo Fisher Scientific), operated at 1.18 kV. A section of 50 nm thickness was removed after each imaging from the imaged block face. The acquired SEM images were aligned and the chloroplast membrane was manually outlined and segmented using the Amira (Thermo Scientific) software.

## Authors’ Contribution

Irem Yilmazer: Conceptualization, Investigation, Methodology, Validation, Formal analysis, Data Curation, Writing – review and editing, Visualization

Pamela Vetrano: Conceptualization, Investigation, Methodology, Validation, Formal analysis, Data Curation, Writing – review and editing, Visualization

Simona Eicke: Investigation, Methodology, Validation, Formal analysis, Data Curation, Visualization

Melanie R. Abt: Investigation, Methodology, Validation, Formal analysis, Data Curation

Eleonora Traverso: Investigation, Methodology, Validation, Formal analysis, Data Curation and Visualization

Tomas Morosinotto: Methodology, Validation, Data Curation, Formal analysis, Visualization, Supervision and Funding Acquisition

Samuel C. Zeeman: Validation, Data Curation, Visualization, Supervision, Project administration, Funding Acquisition

Silvia Ramundo: Conceptualization, Investigation, Methodology, Validation, Resources, Data Curation, Writing – original draft preparation, Writing – review and editing, Visualization, Supervision, Project administration, Funding Acquisition

Mayank Sharma: Conceptualization, Investigation, Methodology, Validation, Resources, Data Curation, Writing – original draft preparation, Writing – review and editing, Visualization, Supervision, Project administration, Funding Acquisition

## Acknowledgements

MS would like to acknowledge the financial support of ETH Zurich Career Seed Award (SEED-29 21-2). MS, SCZ, and SE would like to acknowledge the proteomics/protein analysis service group of Functional Genomic Center Zurich and Dr. Tobias Kockmann for their help in mass spectrometry and proteomics data analysis of Arabidopsis IP samples, the Scientific Center for Optical and Electron Microscopy (ScopeM, ETH Zurich), and the Center for Microscopy and Image Analysis (University of Zurich) for providing access to microscopy facilities. SR and PV acknowledge the financial support of the Austrian Academy of Science and are grateful to the entire team at the Protein Chemistry Facility at IMP/IMBA/GMI for carrying out the mass spectrometry and proteomics data analysis of the *C. reinhardtii* IP samples utilizing the Vienna BioCenter Core Facilities (VBCF) instrument pool. They also wish to thank Nicole Drexter at VBCF for her assistance in imaging *C. reinhardtii* strains through electron microscopy and Sven Klumpe, Ben Engel, Michael Schroda and all Ramundo lab members for discussions on this project. Authors would like to thank Natalie Page and J. Matthew Watson for editing and proofreading, and Liam Dolan and Youssef Belkhadir for critical comments on this manuscript.

