## Supplementary material for "A conserved ESCRT-II-like protein participates in the biogenesis and maintenance of thylakoid membranes": Suppl. File 1

### Supplementary File 1

#### A conserved ESCRT-II-like protein participates in the biogenesis and maintenance of thylakoid membranes

Irem Yilmazer[1#], Pamela Vetrano[2#], Simona Eicke[1], Melanie R. Abt[1], Eleonora Traverso[3], Tomas Morosinotto[3], Samuel C. Zeeman[1], Silvia Ramundo[2*], Mayank Sharma[1*]

1-Institute of Molecular Plant Biology, Department of Biology, ETH Zürich, Zürich, Switzerland
2-Gregor Mendel Institute of Molecular Plant Biology, Vienna Biocenter, Vienna, Austria
3-Department of Biology, University of Padova, Padova, Italy

This file contains Figures S1-S12, Link to Video S1-S2 and Tables S1-S3

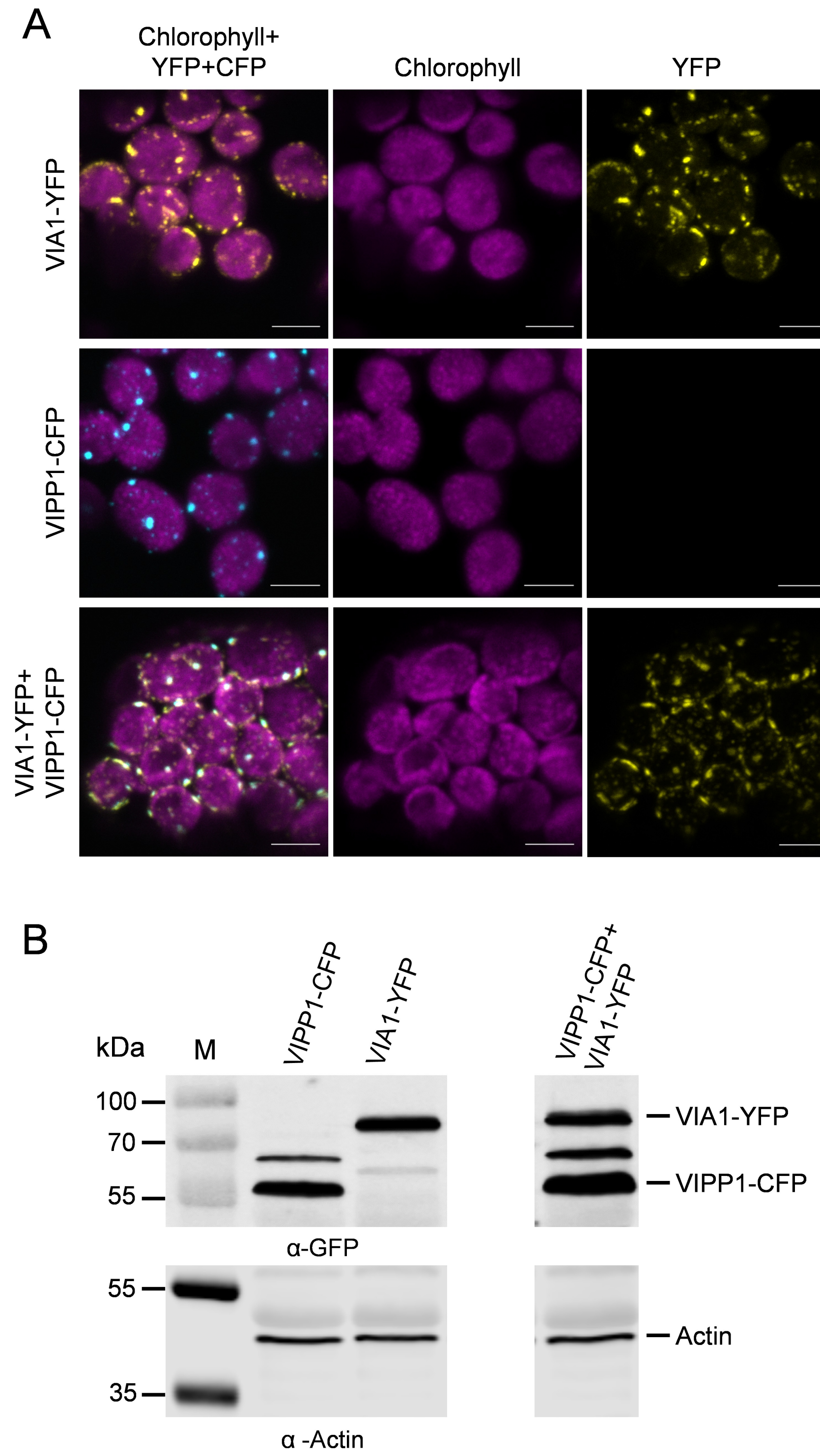

**Figure S1**. **Subcellular Localization of *At*VIA1 and AtVIPP1 in Mesophyll Chloroplasts.** **(A)** Confocal laser scanning microscopy analysis of chloroplasts in mesophyll cell of *N. benthamiana* leaf tissue transiently expressing VIA1-YFP, VIPP1-CFP, or both. Chlorophyll signals are false colored magenta, YFP signals yellow, and CFP signals cyan. Scale bar represents 5 µm. **(B)** Immunoblots against total proteins from leaf tissue expressing the indicated proteins. Similar sized leaf discs were harvested, homogenized in 1X Laemmli buffer, boiled for 5 min at 95°C, and equal volumes of the supernatant were loaded in on 10% SDS-PAGE gels. Proteins were transferred to PVDF membrane and detected with anti-GFP (rabbit) and anti-Actin (mouse) antibodies.

| Entry | Chain | RMSD | TM-score | Identity | Equivalent Residues | Sequence Length | Modelled Residues |
|---|---|---|---|---|---|---|---|
| *At*VIA1 | A | - | - | - | - | 533 | 533 |
| *Cr*VIA1 | A | 3.07 | 0.72 | 38% | 424 | 459 | 459 |

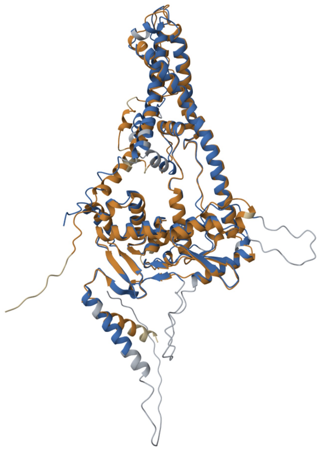

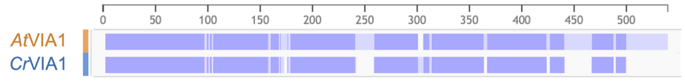

**Figure S2**. **Pairwise Structure Alignment Between *At*VIA1 and *Cr*VIA1.**
Structural alignment between *A. thaliana* VIA1 (in orange) and *C. reinhardtii* VIA1 (in blue) was performed using Pairwise Structure Alignment from RCSB-PDB (1). The alignment analysis includes the root mean square deviation score, a widely used metric for comparing two proteins, the template modeling score (2), which measures topological similarity, and the percentage of sequence identity.

1. H. Berman, K. Henrick, G. Kleywegt, H. Nakamura, J. Markley, RCSB PDB (2012).

2. J. Xu, Y. Zhang, How significant is a protein structure similarity with TM-score = 0.5? *Bioinformatics* **26**, 889–895 (2010).

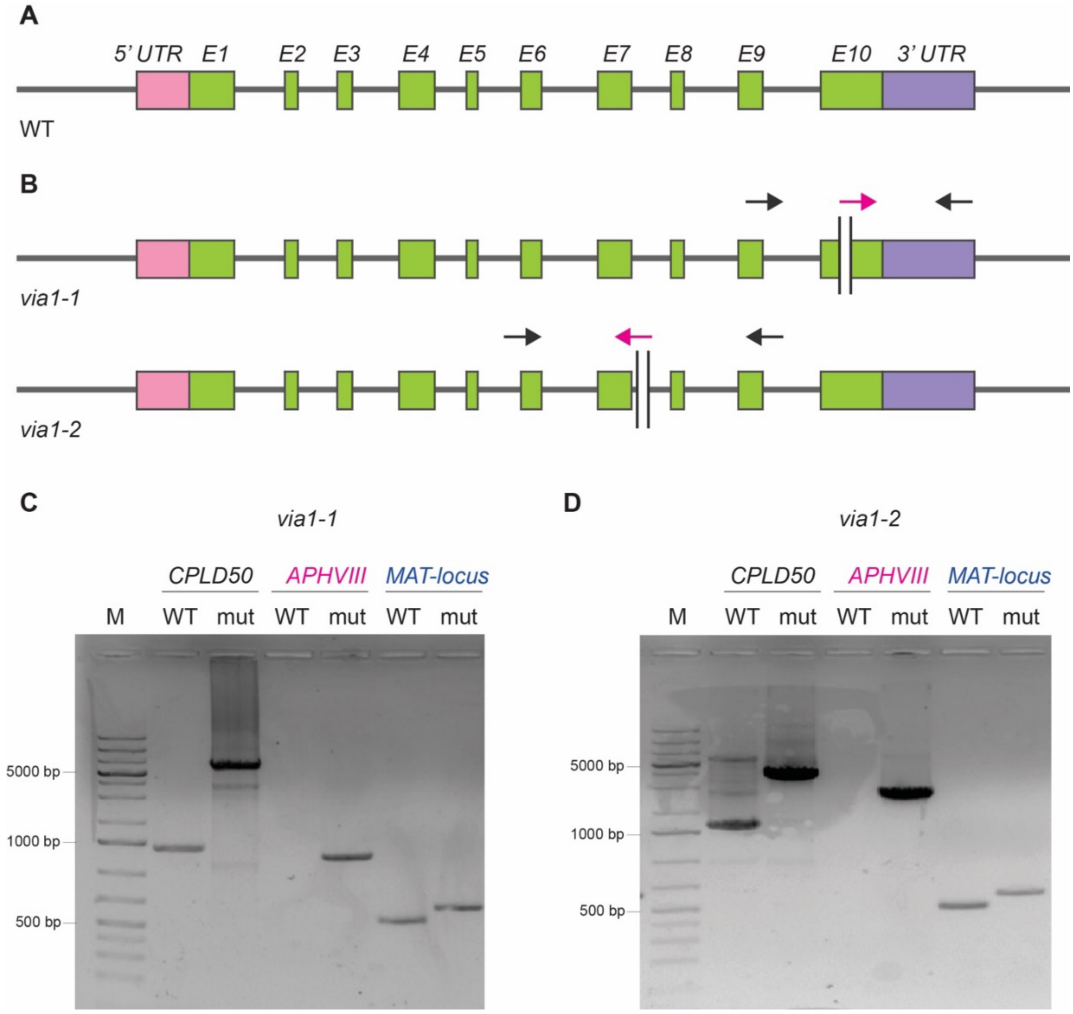

**Figure S3**. **Validation of Mutagenic Insertion in Chlamydomonas *via1-1* and *via1-2* by PCR.**

(A) Schematic representation of the *VIA1* transcript: exons are shown in green, 5'UTR in pink, and 3'UTR in purple. (B) Diagram illustrating the mutagenic insertion carrying the gene conferring paromomycin resistance (*APHVIII*) in the *VIA1* transcript of *via-1* and *via1-2*. The mutagenic cassette (represented by black vertical lines) is located in exon ten and intron seven, respectively. Its orientation is 5'-3' in *via1-1* and 3'-5' in *via1-2*. Arrows indicate primers used for the PCRs shown in panels C and D. (C-D) PCRs validating the mutagenic insertion in *via1-1* (panel C) and *via1-2* (panel D): The first PCR (*VIA1*) amplifies a region between black arrowheads in the diagrams. The size difference between WT and mutant amplicons reflects the presence of the cassette in mutants. The second PCR (*APHVIII*) amplifies the region between black and pink arrowheads, producing an amplicon only in mutants. The third PCR (*MAT-locus*) is a mating-type specific PCR used as a control to ensure the quality of the employed genomic DNA.

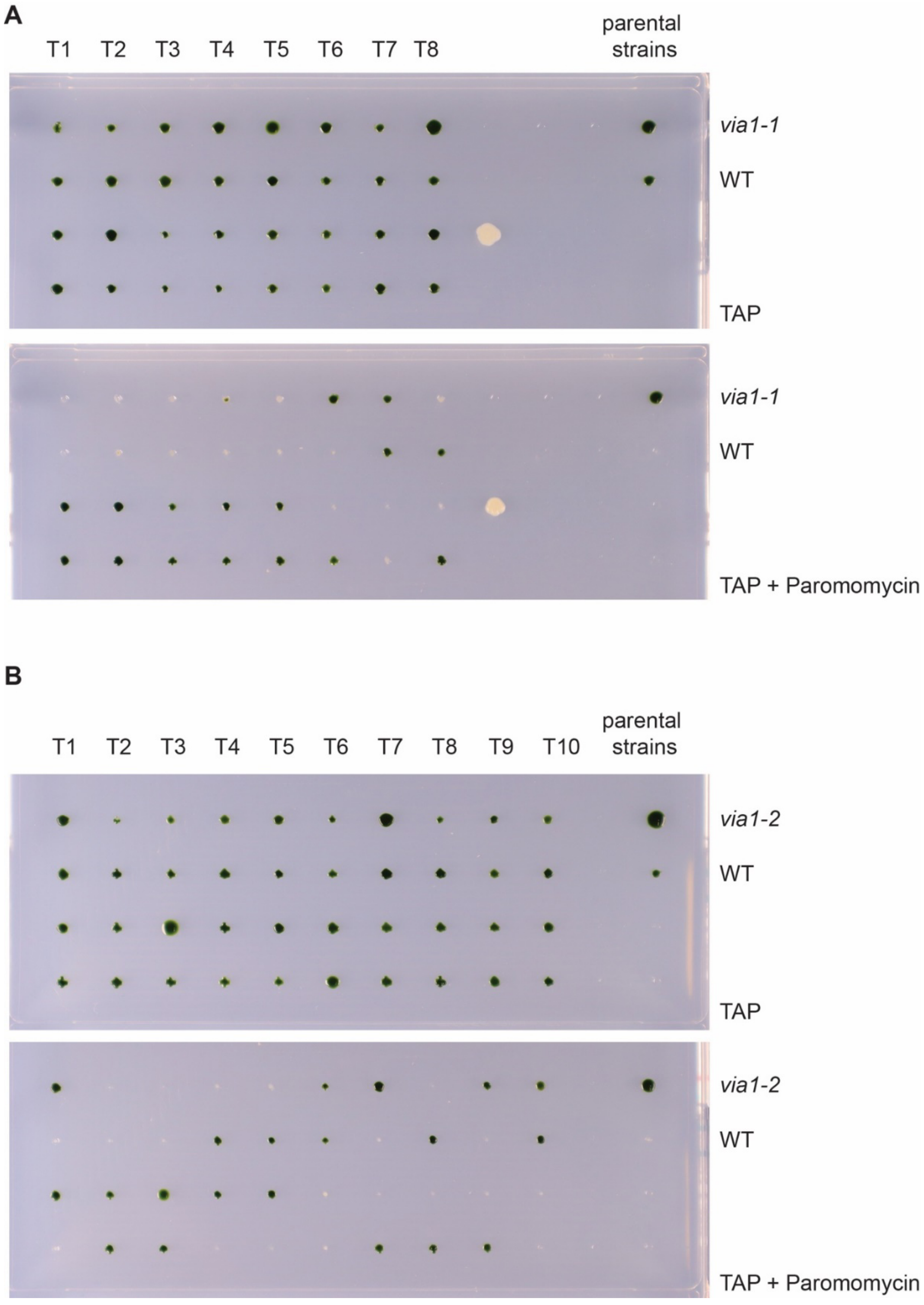

**Figure S4**. **Segregation Analysis of the Mutagenic Cassette in Tetrads from the Backcross of Chlamydomonas *via1-1* or *via1-2* to WT.** Tetrads (T1-T10) derived from the backcross of *via-1* (**A**) or *via1-2* (**B**) to WT strain were re-arrayed on rectangular agar TAP plates. The upper section of each panel shows the tetrads growing on TAP agar, while the lower section displays the tetrads growing on TAP agar supplemented with paromomycin.

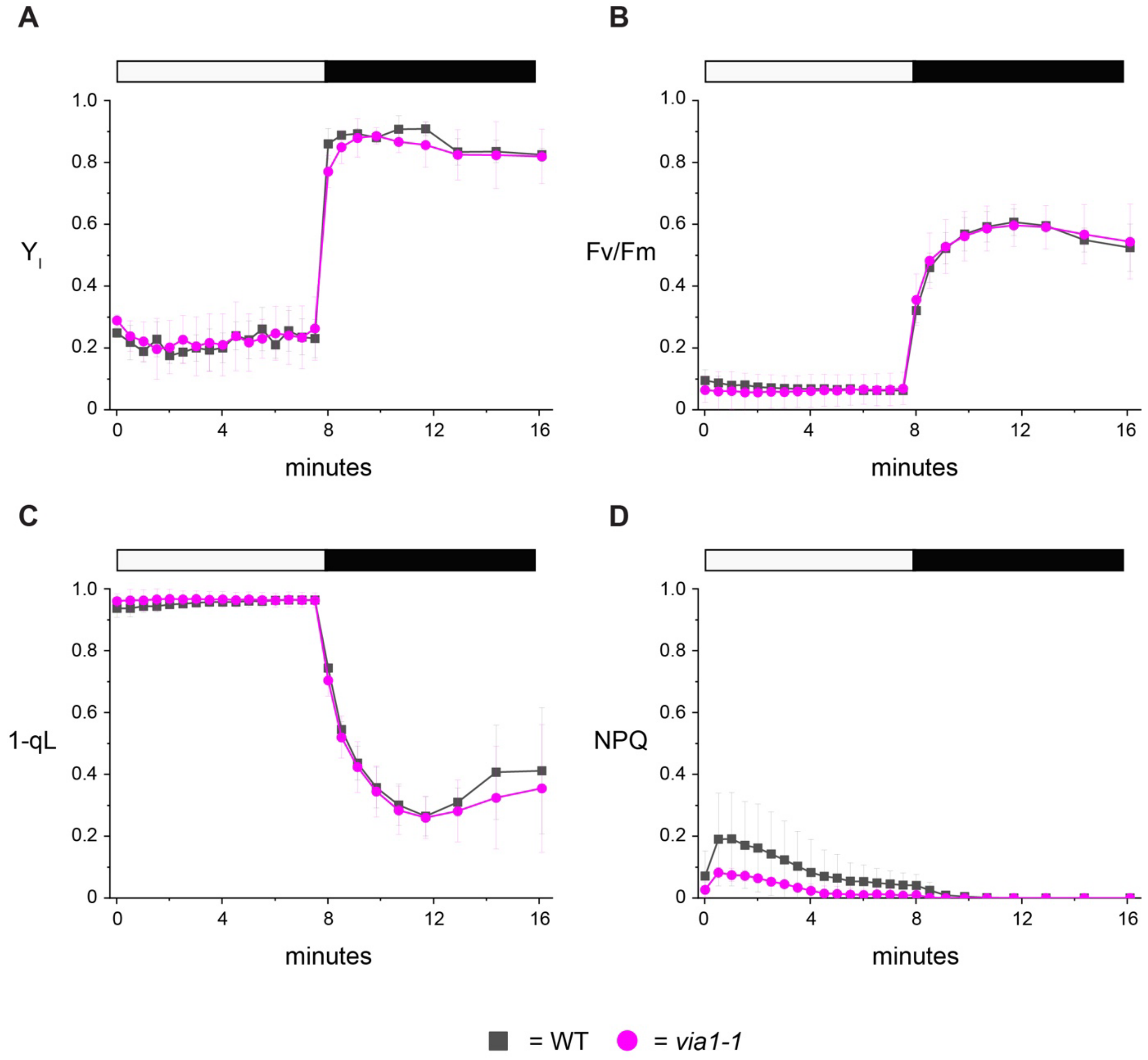

**Figure S5**. **Photosynthetic Measurements in Chlamydomonas WT and *via1-1* Cells Using PAM Fluorometry. (A)** Photosystem I quantum yield (YI) and **(B)** Photosystem II quantum yield (Fv/Fm) were monitored during an 8-minute exposure to actinic light at an intensity of 1000 µmol photons $m^{-2}$ $s^{-1}$, followed by an 8-minute dark period. The Plastoquinone redox state was assessed by analyzing the fluorescence parameters **(C)** 1-qL and **(D)** NPQ. Data are presented as the means ± standard deviation from four independent measurements.

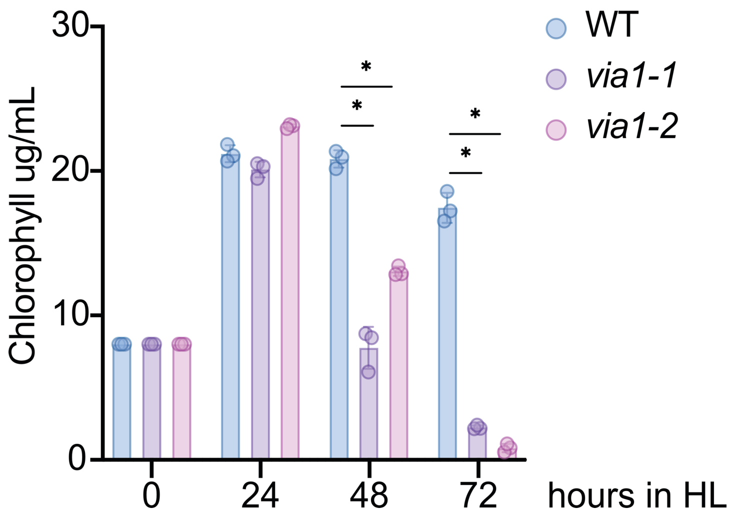

**Figure S6**. **Chlorophyll Measurements in Chlamydomonas WT, *via1-1* and *via1-2* Cells.** Total chlorophyll content in Chlamydomonas strains was measured at various time points during a 3-day high light (HL) treatment. Three replicates were measured for each strain. The high-light (HL) treatment was performed with an intensity of 1000-1200 μmol photons $m^{-2}$ $s^{-1}$.

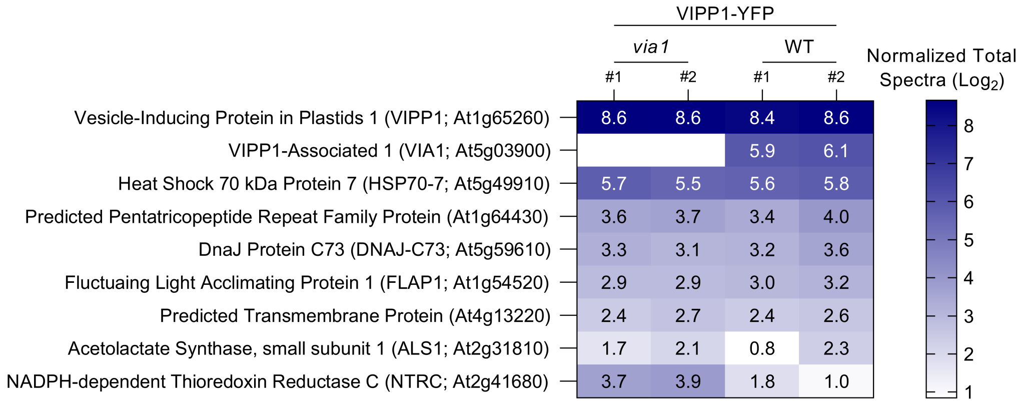

**Figure S7. *At*VIPP1-YFP AP-MS Analysis of Arabidopsis *via1* Plants.** A heatmap representing proteins identified in an AP-MS experiment conducted with Arabidopsis leaf total protein lysates expressing VIPP1-YFP either in *via1* or wild-type background. VIPP1-YFP was isolated using GFP-trap magnetic beads and subjected to mass spectrometry analysis to test for the presence of interacting VIA1 peptides. Scaffold proteomics software was employed to analyze the identified peptides, and a $Log_2$-transformed, quantitative value representing the identified peptide (normalized total spectra) is displayed within each cell of the heatmap. Empty cells represent the absence of corresponding peptides. As anticipated, peptides corresponding to VIA1 were identified in samples obtained from plants expressing VIPP1-YFP in a wild-type background. However, in samples from plants expressing VIPP-YFP in the *via1* background, no VIA1-specific peptides were detected. Notably, the other interacting proteins were present at comparable intensities, indicating that the *via1* mutants lack VIA1 protein.

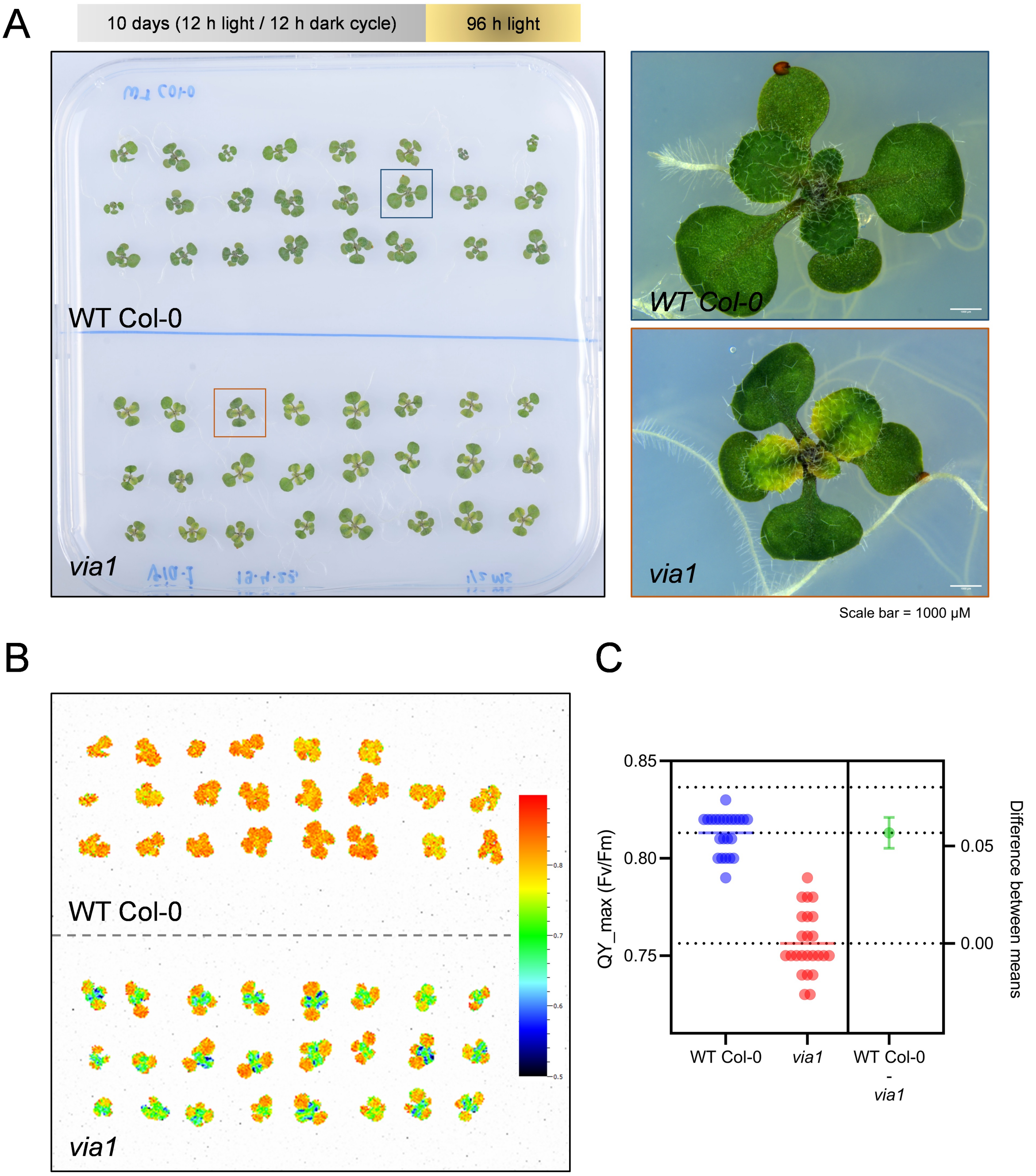

**Figure S8. Photosynthetic Measurements of Arabidopsis WT and *via1* Seedlings. (A)** A photograph of Arabidopsis seedlings corresponding to WT Col-0 and *via1* lines cultivated on ½ strength MS plates for 10 days under a 12 h light, 12 h dark cycle, followed by an additional 96 h under continuous light. **(B)** An image of the same seedlings, captured with a FluorCam, illustrating the maximum quantum yield of Photosystem II (Fv/Fm) values. The Fv/Fm values are represented on a scale from 0.5 (blue) to 0.9 (red). **(C)** Fv/Fm values of individual seedlings from the left panel. Statistical significance was determined using an unpaired t-test with Welch's correction, resulting in a P-value of <0.0001, indicating significant differences between the two measurements. The third column represents the difference between the means of WT Col-0 and *via1* measurements.

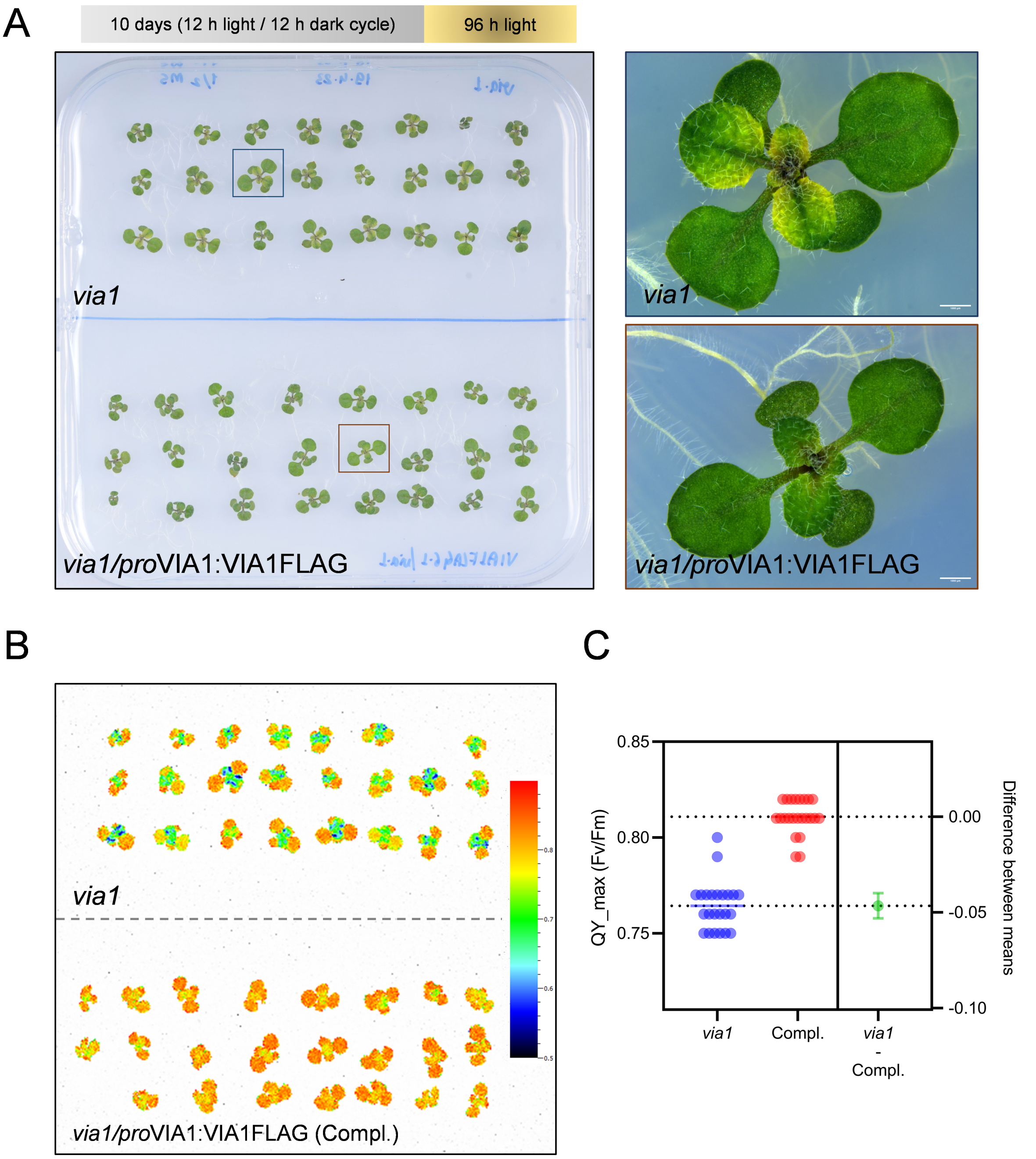

**Figure S9. Photosynthetic Measurements of Arabidopsis *via1* and *via1/pro*VIA1:VIA1FLAG Complemented Seedlings*.* (A)** A photograph of Arabidopsis seedlings corresponding to *via1* and *via1/pro*VIA1:VIA1FLAG complementation lines, cultivated on ½ strength MS plates for 10 days under a 12 h light and 12 h dark cycle, followed by an additional 96 h under continuous light. **(B)** An image of the aforementioned seedlings, captured with a FluorCam, illustrating the maximum quantum yield of Photosystem II (Fv/Fm) values. The Fv/Fm values are represented on a scale from 0.5 (blue) to 0.9 (red). **(C)** Fv/Fm values of individual seedlings from the left panel are plotted. Statistical significance was determined using an unpaired t-test with Welch's correction, resulting in a P-value of <0.0001, indicating significant differences between the two measurements. The third column represents the difference between the means of *via1/pro*VIA1:VIA1FLAG (Compl.) and *via1* measurements.

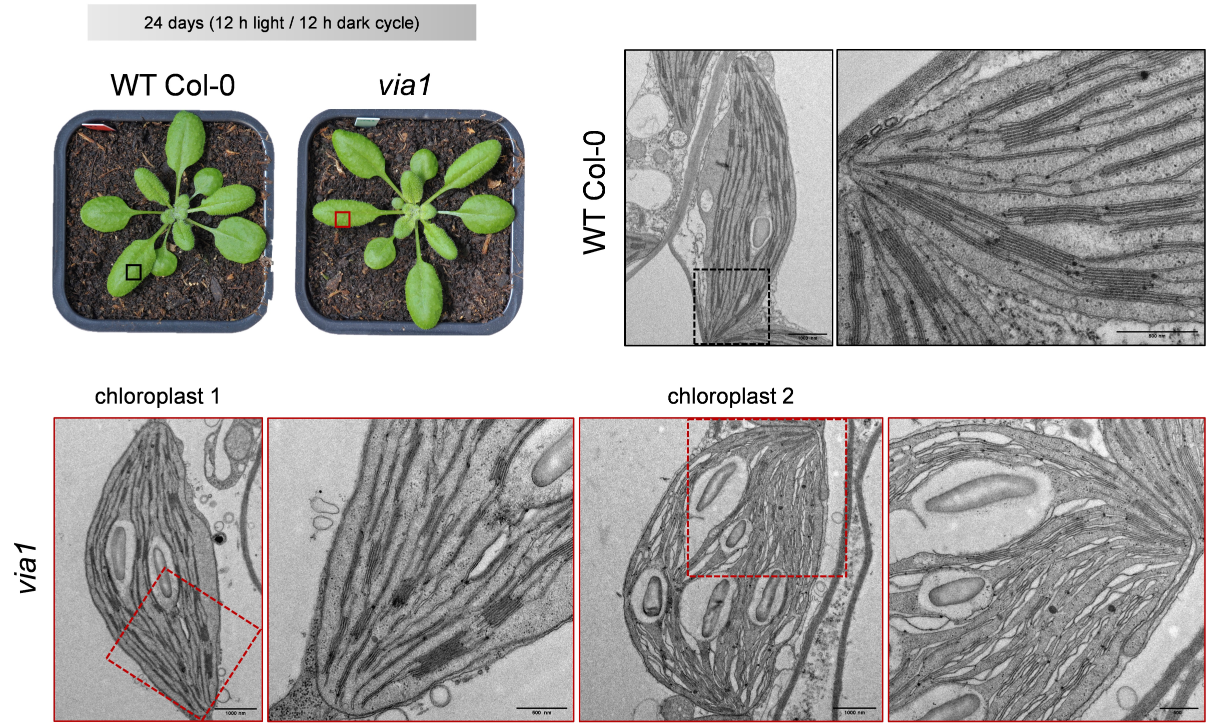

**Figure S10. Transmission Electron Micrographs of Chloroplasts from Mature Leaf Sections of Arabidopsis Wild Type (WT Col-0) and *via1* Plants.** Arabidopsis plants were grown under a 12 h light and 12 h dark growth cycle for 24 days. Tissue sections from the middle-part of mature leaves were harvested after 90 min. of light exposure on 24$^{th}$ day, immediately fixed and embedded in Spurr resin, and imaged. Top panels display a chloroplast from WT plants (black outline) while both chloroplasts in bottom panel belongs to *via1* plants (red outline). Squares in the images represent the area chosen for imaging at higher magnifications. In *via1* leaves, only a few chloroplasts with swollen thylakoids (like chloroplast 2) were observed, while the majority of chloroplasts displayed morphology similar to chloroplast 1. Scale bars corresponds to 1000 nm for the full chloroplast images and 500 nm for the higher magnification images.

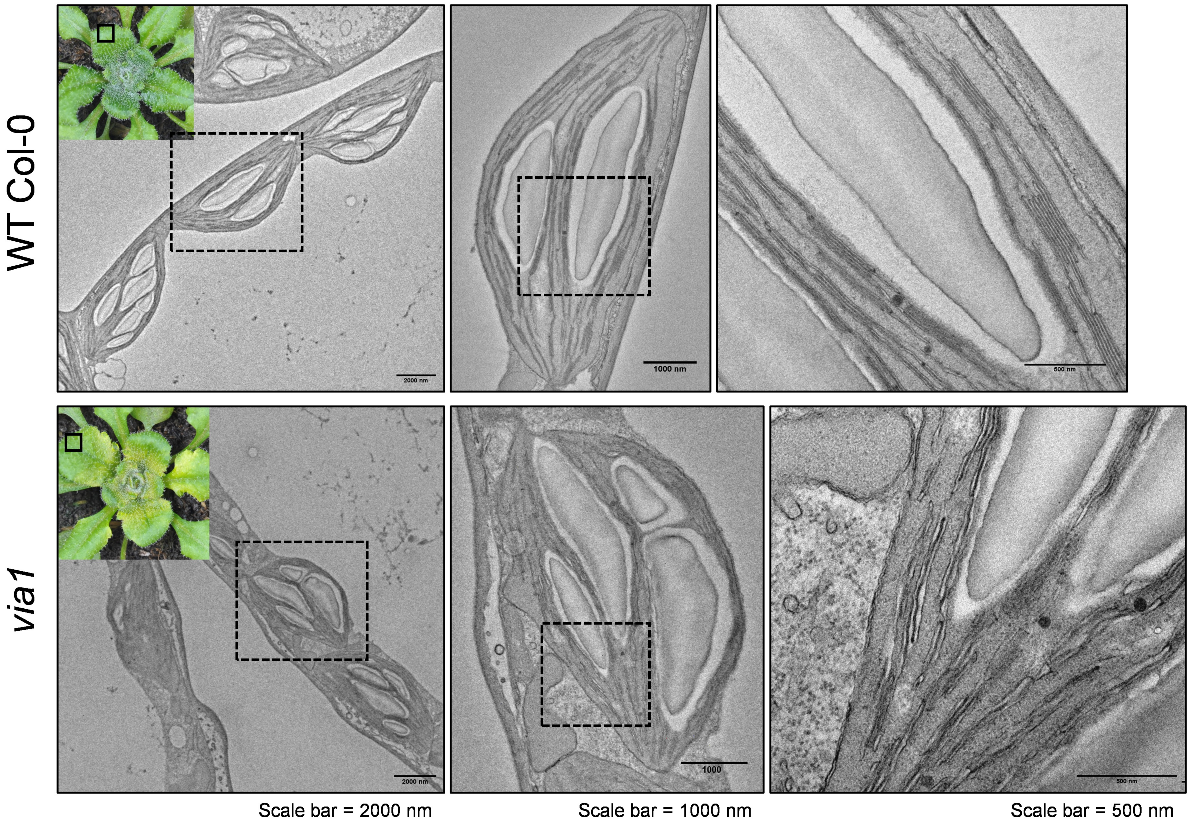

**Figure S11. Transmission Electron Micrographs of Chloroplasts from the Green Region of the Leaf Tissue of Arabidopsis Wild Type (WT Col-0) and *via1* Plants.** Arabidopsis plants were grown on soil under a 12 h light and 12 h dark growth cycle for 24 days, followed by 96 h of continuous light exposure. Green segments from the newly emerging leaves of corresponding lines were harvested, immediately fixed, embedded in Spurr resin and imaged. The material was collected from the same plants used for generating images in Figure 9A.

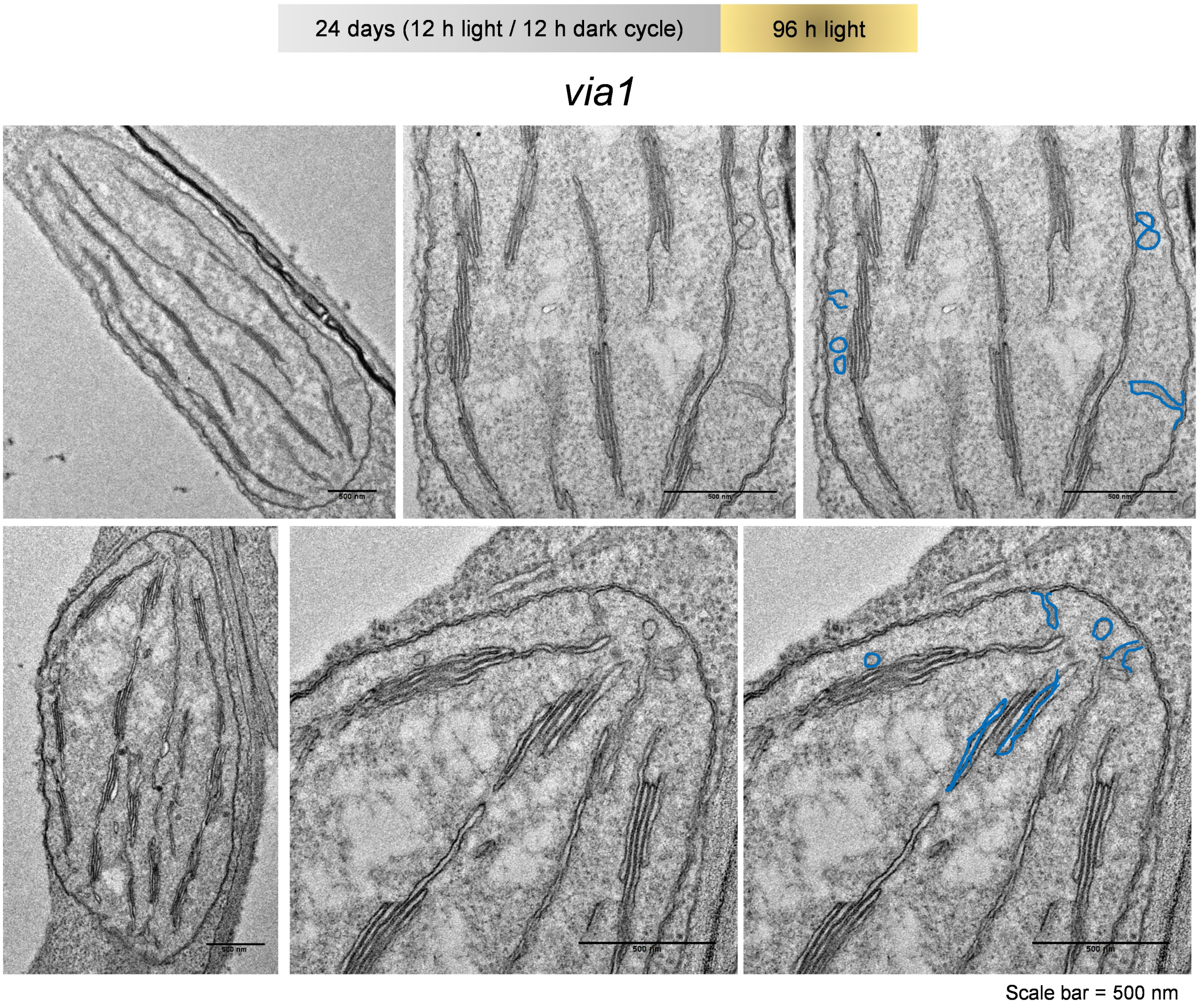

**Figure S12. Transmission Electron Micrographs of Chloroplasts from the Base of Newly Emerging Pale Green Leaf Tissue of Arabidopsis *via1* Plants**. Arabidopsis plants were cultivated in soil under a 12 h light and 12 h dark growth cycle for 24 days, followed by 96 h of continuous light exposure. Leaf tissue sections from the central pale green regions of the rosette were collected, immediately fixed, embedded, and imaged using a transmission electron microscope. The resin blocks used for imaging were the same as those used to generate Figure 9A. These chloroplasts exhibit a significant reduction in the thylakoid network and the accumulation of membrane invaginations and vesicular structures. In the right panel, the vesicles and membrane invaginations have been manually highlighted in blue.

**Video S1.** https://polybox.ethz.ch/index.php/s/bxzmYLxyjeuIcMt
A 3D reconstruction and segmentation of SEM images stack highlighting the chloroplasts from WT Col-0 leaf tissue exposed to continuous light (related to Figure 9B).

**Video S2.** https://polybox.ethz.ch/index.php/s/mGygzhbnFeuJgfo
A 3D reconstruction and segmentation of SEM images stack highlighting the chloroplasts from *via1* leaf tissue exposed to continuous light (related to Figure 9B).

**Table S1.** List of Oligonucleotides Used in This Study.

| Primer name | Sequence 5' → 3' |
|---|---|
| oRS01 | CCGAGGAGAAACTGGCCTT |
| oRS30 | GCACACGCTAACATCTACAA |
| oRS37 | GCTTCGTGGAGTCCATCTTC |
| oRS38 | CCTAGTCCACCCTCACCGTA |
| oRS39 | TACGGTGAGGGTGGACTAGG |
| oRS40 | GGTATGAAACCTCTCGCCAA |
| oRS47 (qRT_*A*_Fw) | TGTTCATGGACGTGACGGAC |
| oRS48 (qRT_*A*_Rv) | CCCTTGGCTCGGATCATCTC |
| oRS57 (qRT_*B*_Fw) | GTCTGTGTGTGTTGTGCAATGGG |
| oRS58 (qRT_*B*_Rv) | GCTCTCTCTCGTGTCACCTTTG |
| oRS59 (qRT_*C*_Fw) | ATCTTGTGGACCGGCCTTTT |
| oRS60 (qRT_*C*_Rv) | GTCGTACACCACGTCATCGT |
| oRS61 (qRT_*D*_Fw) | GACGATGACGTGGTGTACGA |
| oRS62 (qRT_*D*_Rv) | ACTGTCCCAGTCGTCCAGAT |
| qRT_GBLP_Fw | CAAGTACACCATTGGCGAGC |
| qRT_GBLP_Rv | CTTGCAGTTGGTCAGGTTCC |
| VIA1_attB1_F | GGGGACAAGTTTGTACAAAAAAGCAGGCTTCACCATGGCGTGTGTATCGACATG |
| VIA1_attB2_R | GGGGACCACTTTGTACAAGAAAGCTGGGTCATCCGATTTTTCTAACTCTTTGAATC |
| VIA1_prom_attB1 | GGGGACAAGTTTGTACAAAAAAGCAGGCTTCACCCAGATAGCTCATATA TACTCAAAAAAC |
| VIPP1_attB1_F | GGGGACAAGTTTGTACAAAAAAGCAGGCTTCACCATGGCTCTCAAAGCTTCACCT GTTAC |
| VIPP1_attB2_R | GGGGACCACTTTGTACAAGAAAGCTGGGTCAAAGTCGTTAGCTTTCCTTCGCAG |
| cpFeS_attB1_F | GGGGACAAGTTTGTACAAAAAAGCAGGCTTCACCATGGCTTTCGCTACTGGAATC |
| cpFeS_attB2_R | GGGACCACTTTGTACAAGAAAGCTGGGTCCATCTCGGCAGCAAAAGACTTC |
| VIA1_RT_5'_F | CGATGGATGCGGTAGATGAGTGTG |
| VIA1_RT_5'_R | GCTCGTCGCCTGTTGTAATAATTGG |
| VIA1_RT_3'_F | TCCCTTTGATTCGGTGGTTTTCAA |
| VIA1_RT_3'_R | CCAGCGTCCATGAATGTTTTTACG |
| Actin_RT_F | TCCTCTCCGCTTTGAATTGTCTCG |
| Actin_RT_R | TGATGTCTTGGCCTACCAACAACAC |
| VIA1_LP | TTTGACGATTATCATCCTCGC |
| VIA1_RP | GATTTTGCTGCTCTATCGTCG |
| SALK_Lba1 (BP) | TGGTTCACGTAGTGGGCCATCG |

**Table S2.** List of Reagents Used in This Study.

| Reagent type | Designation | Source | Identifiers | Additional information |
|---|---|---|---|---|
| Gene (*C. reinhardtii*) | VIPP1 | Phytozome | Cre13.g583550 | https://phytozome-next.jgi.doe.gov |
| Gene (*C. reinhardtii*) | VIA1, CPLD50, VPL3 | Phytozome | Cre07.g338350 | https://phytozome-next.jgi.doe.gov |
| Gene (*A. thaliana*) | VIPP1 | TAIR | At1g65260 | www.arabidopsis.org |
| Gene (*A. thaliana*) | VIA1 | TAIR | At5g03900 | www.arabidopsis.org |
| Gene (*A. thaliana*) | cpFeS (CPISCA) | TAIR | At1g10500 | www.arabidopsis.org |
| Antibody | anti-VIPP1 (rabbit, polyclonal) | Gift from Prof. Jean-David Rochaix | | (1:20000) |
| Antibody | anti-α-tubulin (mouse, monoclonal) | Sigma | T6074 | (1:10000) |
| Antibody | anti-ClpP1 (rabbit, polyclonal) | Gift from Dr. Olivier Vallon | | (1:5000) |
| Antibody | anti-PsaA (rabbit, polyclonal) | Gift from Prof. Jean-David Rochaix | | (1:10000) |
| Antibody | anti-D1 (rabbit, polyclonal) | Gift from Prof. Jean-David Rochaix | | (1:3000) |
| Antibody | anti-D2 (rabbit, polyclonal) | Gift from Prof. Jean-David Rochaix | | (1:3000) |
| Antibody | anti-cytF (rabbit, polyclonal) | Gift from Prof. Jean-David Rochaix | | (1:10000) |
| Antibody | HRP-conjugated anti-rabbit | Promega | W4011 | (1:10000) |
| Antibody | HRP-conjugated anti-mouse | Promega | W4021 | (1:10000) |
| Antibody | Anti-GFP (rabbit, polyclonal) | Torrey Pines Biolabs | TP401 | (1:10000) |
| Antibody | anti-FLAG (mouse, monoclonal) | Sigma Aldrich | F1804 | (1:1000) |
| Antibody | IRDye® 800CW (donkey anti-rabbit IgG) | LI-COR | 926-32213 | (1:10000) |
| Antibody | IRDye® 680RD (goat anti-mouse IgG) | LI-COR | 926-68072 | (1:10000) |
| Antibody | Anti-Actin (rabbit, polyclonal) | Sigma Aldrich | A0480 | (1:10000) |
| Commercial kit | KOD Hot Start DNA Polymerase | EMD Millipore | 71086-5 | |
| Commercial kit | Phusion High-Fidelity DNA Polymerase | ThermoFisher Scientific | F530L | |
| Commercial kit | Direct-zol DNA Miniprep Plus | Zymo Research | D4019 | |
| Commercial kit | Zymoclean Gel DNA Recovery Kit | Zymo Research | D4001 | |

| | | | |
|---|---|---|---|
| Commercial kit | In-Fusion HD cloning plus | Takara | 38910 |
| Commercial kit | SuperSignal West Femto | ThermoFisher Scientific | 34095 |
| Commercial kit | SuperSignal West Dura | ThermoFisher Scientific | 34075 |
| Commercial kit | Protease Inhibitors EDTA-free | Roche | 11873580001 |
| Commercial kit | Dynabeads™ Protein A | Invitrogen | 10002D |
| Commercial kit | BP Clonase-II | ThermoFisher Scientific | 11789020 |
| Commercial kit | LR Clonase-II | ThermoFisher Scientific | 11791020 |
| Commercial kit | GFP-Trap® Magnetic Agarose | Chromtek | gtma |
| Commercial kit | RevertAid First Strand cDNA Synthesis Kit | ThermoFisher Scientific | K1621 |
| Chemical compound, drug | Bicinchoninic Acid solution | Sigma | B9643-1L |
| Chemical compound, drug | Digitonin | EMD Millipore | 300410-5GM |
| Chemical compound, drug | Spectinomycin | Sigma | S4014-5G |
| Chemical compound, drug | Lugol’s solution | Gatt-Koller | 403096459 |
| Chemical compound, drug | Maxima First Strand cDNA | ThermoFisher Scientific | K1641 |
| Chemical compound, drug | Fast Start Essential DNA Green Master | Roche | 06402712001 |
| Equipment | MagnaRack | Invitrogen | CS15000 |
| Equipment | Kontes Duall #21 homogeniser | Kimble | 8854500021 |
| Equipment | Mixer mill MM400, grinding jars (50ml) and grinding balls (20mm) | Retsch | 70354 |
| Equipment | Dissection microscope (SporePlay+) | Singer | SPO-002 |
| Equipment | Magnetic stand MagnaRack | Invitrogen | CS15000 |
| Equipment | Plant LED Grow Light | Phlizon | Cob Series 3000W |
| Equipment | Imaging system | iBright CL1500 | A44114 |
| Equipment | Nitrocellulose Blotting Membrane | Amersham | 10600001 |
| Equipment | Scalpel | Aesculap AG | BA821SU |
| Equipment | Razor blade | Apollo Herkenrath | 10-210-090 |
| Equipment | SureBeads™ Magnetic Rack | BioRad | 1614916 |
| Software | Sequence Data Analysis | Snapgene | |
| Software | Statistic Data Analysis | Graphpad | |

| | | | | |
|---|---|---|---|---|
| Software | Molecular Visualization Program | ChimeraX | | |
| Software | Protein Structure Prediction | Alphafold Google CoLab | | |
| Software | Protein structure alignment | RCSB PDB | | https://www.rcsb.org/alignment |
| Software | DeepTMHMM | Technical University of Denmark | | https://dtu.biolib.com/DeepTMHMM/ |
| Software | Phylogenetic Tree | MEGA 7 | | |
| Software | Image Segmentation | Amira | | |
| Software | Image Processing | Fiji | | |
| Software | DNA Sequence Analysis | CLC Genomics | | |

**Table S3.** List of Biological Resources.

| Identifier | Strain name | Description | Comments | Reference |
|---|---|---|---|---|
| CrRS166 | CC-5325 | This wild-type strain is a haploid progeny from a cross between 4A (mt-) (Dent et al. 2005) and D66 (mt+) (Schnell and Lefebvre, 1993) | This strain was employed for all the experiments except for the backcross and the qRT-PCRs. | Li et al, The Plant Cell, 2016<br><br>(Available at the Chlamydomonas Resource Center |
| CrRS197 | XF305 F1 | This wild-type strain is a haploid progeny from a cross between CC5235 and CC125 (Pröschold T, Harris EH, Coleman AW (2005) | This strain was only employed for the backcross experiment and the qRT-PCRs | Strain generated during this study |
| CrRS432 | *via1-1* | Cre07.g338350 mutant allele<br><br>Clip ID: LMJ.RY0402.176599 | | Li et al, The Plant Cell, 2016<br><br>(Available at the Chlamydomonas Resource Center) |
| CrRS433 | *via1-2* | Cre07.g338350 mutant allele<br><br>Clip ID: LMJ.RY0402.193812 | | Li et al, The Plant Cell, 2016<br><br>(Available at the Chlamydomonas Resource Center) |
| SPV54.1 | T1 to T8 | Plate containing tetrads obtained from backcross of CrRS432 to CrRS197 | | Strain generated during this study |
| SPV54.2 | T1 to T10 | Plate containing tetrads obtained from backcross of CrSR433 to CrRS197 | | Strain generated during this study |
| SALK_057879C | *via1* | A T-DNA insertional line obtained from NASC. | | www.arabidopsis.org |
| VIA1FLAG | *via1*/*pro*VIA1:VIA1FLAG | Homozygous T3 complementation lines, selected based on segregation analysis on hygromycin plates | | Lines generated during this study |
| VIPP1-YFP | VIPP1-YFP-1, VIPP1-YFP-2, VIPP1-YFP-3 | Independent transgenic lines expressing VIPP1-YFP in WT Col-0 background. | T2 segregating population | Lines generated during this study |
| *via1*/ VIPP1-YFP | *via1*/ VIPP1-YFP-1, *via1*/ VIPP1-YFP-2 | Independent transgenic lines expressing VIPP1-YFP in *via1* background. | T2 segregating population | Lines generated during this study |
