## Supplementary figures and images for "A conserved ESCRT-II-like protein participates in the biogenesis and maintenance of thylakoid membranes"

### Suppl. File 3

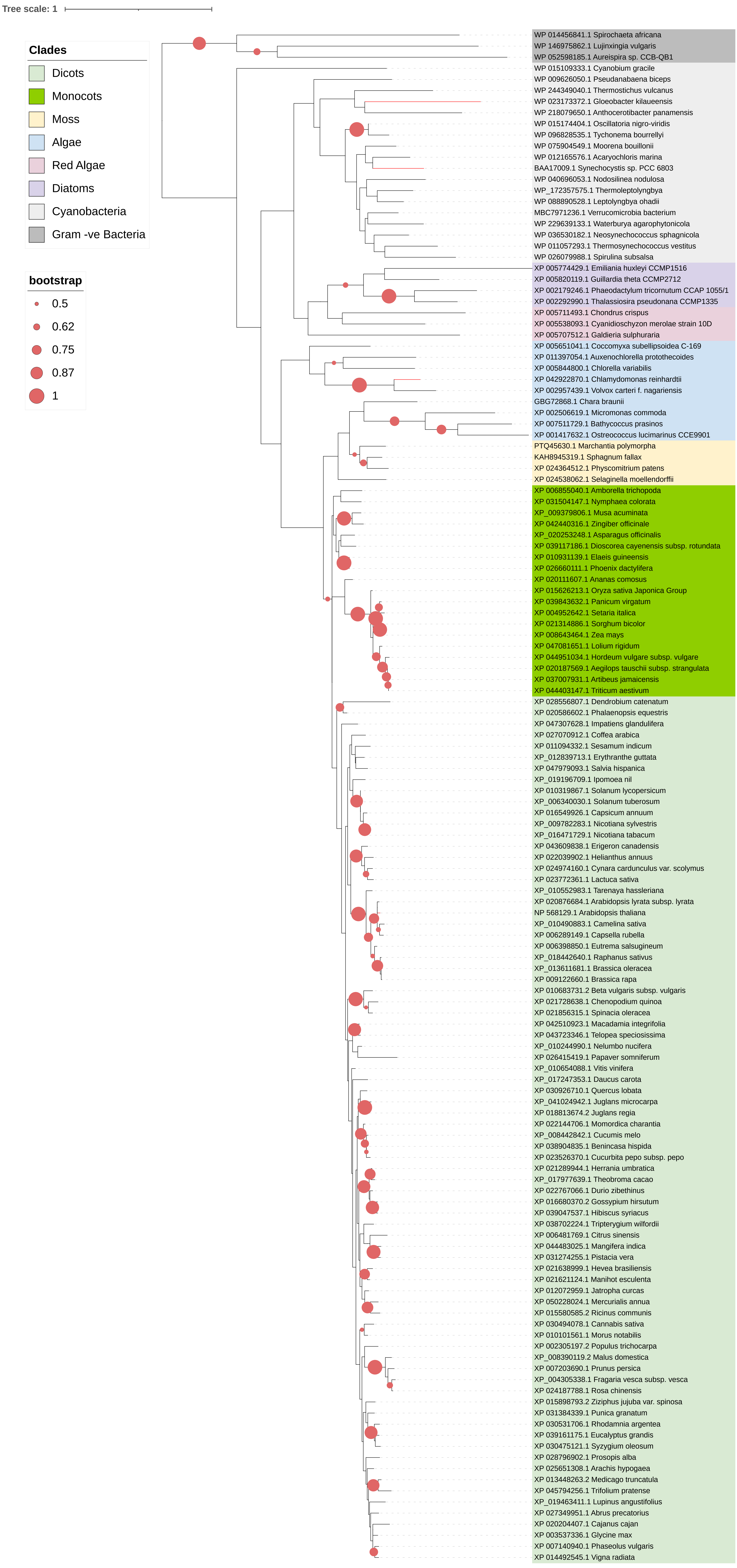
